# Structural venomics: evolution of a complex chemical arsenal by massive duplication and neofunctionalization of a single ancestral fold

**DOI:** 10.1101/485722

**Authors:** Sandy S. Pineda, Yanni K-Y. Chin, Eivind A.B. Undheim, Sebastian Senff, Mehdi Mobli, Claire Dauly, Pierre Escoubas, Graham M. Nicholson, Quentin Kaas, John S. Mattick, Glenn F. King

## Abstract

Spiders are the most successful venomous animals on the planet, with more than 47,000 extant species. Most spider venoms are dominated by disulfide-rich peptides (DRPs) with a diverse range of pharmacological activities. Although some venoms contain thousands of unique peptides, little is known about the mechanisms used to generate such complex chemical arsenals. We used a combined transcriptomic, proteomic and structural biology approach to demonstrate that the lethal Australian funnel-web spider produces 33 superfamilies of venom peptides and proteins, more than described for any other arachnid. We show that 15 of the 26 DRP superfamilies form an ultra-stable inhibitor cystine knot motif, and that these DRPs are the major contributor to the diversity of the venom peptidome. NMR data reveal that most of these DRPs are structurally related and range in complexity from simple to highly elaborated knottin domains that likely evolved from a single ancestral fold.

---

Excluding insects and mites, spiders are the most speciose animals on Earth^1^. Molecular paleobiological analyses indicate that spiders evolved from an arachnid ancestor in the late Ordovician around 450 million years ago^2^, and consequently their venoms are the product of hundreds of millions of years of evolutionary fine-tuning that has produced a complex cocktail of bioactive compounds with a diverse range of pharmacological properties.

Spider venoms are a heterogeneous mixture of salts, low molecular weight organic compounds (<1 kDa), linear and disulfide-rich peptides (DRPs; typically 3–9 kDa with 3–6 disulfide bonds), and proteins (10– 120 kDa)^3–5^. However, peptides are major components of most spider venoms, with some reported to contain more than 1000 peptides^6^. The majority of these peptides are DRPs with masses of 3.0–4.5 kDa (ca. 25–40 residues), although there is a significant fraction with mass of 6.5–8.5 kDa (ca. 58–76 residues)^5^. As the primary function of spider venom is to immobilize prey, it is perhaps not surprising that most spider-venom DRPs that have been functionally characterized target neuronal ion channels and receptors^5,7,8^.

Although spider-venom DRPs have been shown to adopt a variety of three-dimensional (3D) scaffolds, including the Kunitz-type fold^9^, prokineticin/colipase fold^10^, disulfide-directed β-hairpin (DDH)^11^ and inhibitor cystine knot (ICK) motif^12^, the vast majority of spider-venom DRP structures solved to date conform to the ICK motif. The ICK motif is defined as an antiparallel β sheet stabilized by a cystine knot^12^. In spider toxins, the β sheet typically comprises only two β strands, although a third N-terminal strand is present in some cases^13^. The cystine knot comprises a “ring” formed by two disulfide bonds and the intervening sections of the polypeptide backbone with a third disulfide piercing the ring to create a pseudo-knot^12^. The pseudo-knot provides ICK peptides (also known as knottins^14^) with exceptional resistance to chemicals, heat and proteases,^15,16^ which has made them of interest as drug and insecticide leads^5,15,17^. Numerous spider toxins show minor^18^ or more significant^19^ elaborations of the basic ICK fold that involve an additional stabilizing disulfide bond. More recently, “double-knot” spider toxins have been reported in which two structurally independent ICK domains are joined by a short, flexible linker^20–22^.

Like other peptide folds with stabilizing disulfide bridges, knottin DRPs display a remarkable diversity of biological functions including modulation of many different types of ligand-and voltage-gated ion channels^5^. Despite strong conservation of the knottin scaffold across a taxonomically diverse range of spiders, several factors have hampered analysis of their evolutionary history^23^. First, it is not until recently that a large number of knottin precursor sequences have become available from venom-gland cDNA sequencing projects. Second, the disulfide framework in small DRPs generally constrain the peptide fold to such an extent that most non-cysteine residues can be mutated without damaging the peptide’s structural integrity, which is a luxury not afforded to most globular proteins^23^. Thus, evolution of DRPs is typically characterized by the accumulation of many mutations, leaving very few conserved residues available for deep evolutionary analyses^23,24^. Third, very few structures have been solved for spider-venom DRPs larger than 5 kDa, making it unclear whether just a few or many of the larger DRPs are highly derived or duplicated knottins.

The only attempt so far to study the phylogeny of spider-venom knottins concluded that both orthologous diversification and lineage-specific paralogous diversification were important in generating the diverse arsenal of spider-venom knottins^25^. These authors also reported that knottin neurotoxins that act as gating modifiers via binding to the voltage sensor of voltage-gated ion channels likely evolved on multiple independent occasions. However, whether this resulted from convergent functional radiation of toxins or independent evolution of gating modifiers from non-venom knottins remains unknown. The lack of non-venom gland homologues and the use of only a single non-spider knottin outgroup precluded any conclusions regarding the toxin, or non-toxin, origins of these convergently evolved spider-venom knottins. Thus, the deep evolutionary history of spider-venom knottins remains enigmatic^25^.

Here we report the combined use of proteomic, transcriptomic, and structural approaches to determine the complete repertoire and evolutionary history of DRPs expressed in the venom of *Hadronyche infensa*, a member of the family of lethal Australian funnel-web spiders. This holistic approach provided the most comprehensive overview to date of the toxin arsenal of a spider and it revealed that the enormously diverse repertoire of DRPs in spider venom is largely derived from a single ancestral knottin fold.

## Results and Discussion

### Mass spectrometry reveals an exceedingly complex spider-venom peptidome

Mass spectrometry (MS) analyses revealed that the venom peptidome of *Hadronyche infensa* is more complex than that reported for any other terrestrial venomous animal. We used two complementary mass spectrometry platforms to examine the distribution of peptide masses in pooled venom from female *H. infensa*. Analysis of crude venom using matrix-assisted laser desorption/ionization tandem time-of-flight (MALDI-TOF/TOF) or Orbitrap MS resulted in 2053 and 1649 distinguishable peptide masses, respectively (Fig. 1A-D). Only 651 masses were detected in both MS data sets, leading to a total of 3051 unique venom peptides (Fig. 1E). This is considerably more than the ~1000 peptides reported to be present in venom of the Australian funnel-web spider *Hadronyche versuta*^6^, although this difference is likely a reflection of recent improvements in MS sensitivity rather than an indication of major differences in venom complexity between these closely related species. We conclude that the venom peptidome of Australian funnel-web spiders is more complex than reported for any other venomous animal with the exception of marine cone snails, which have similarly complex venoms^26^.

**Figure 1:**
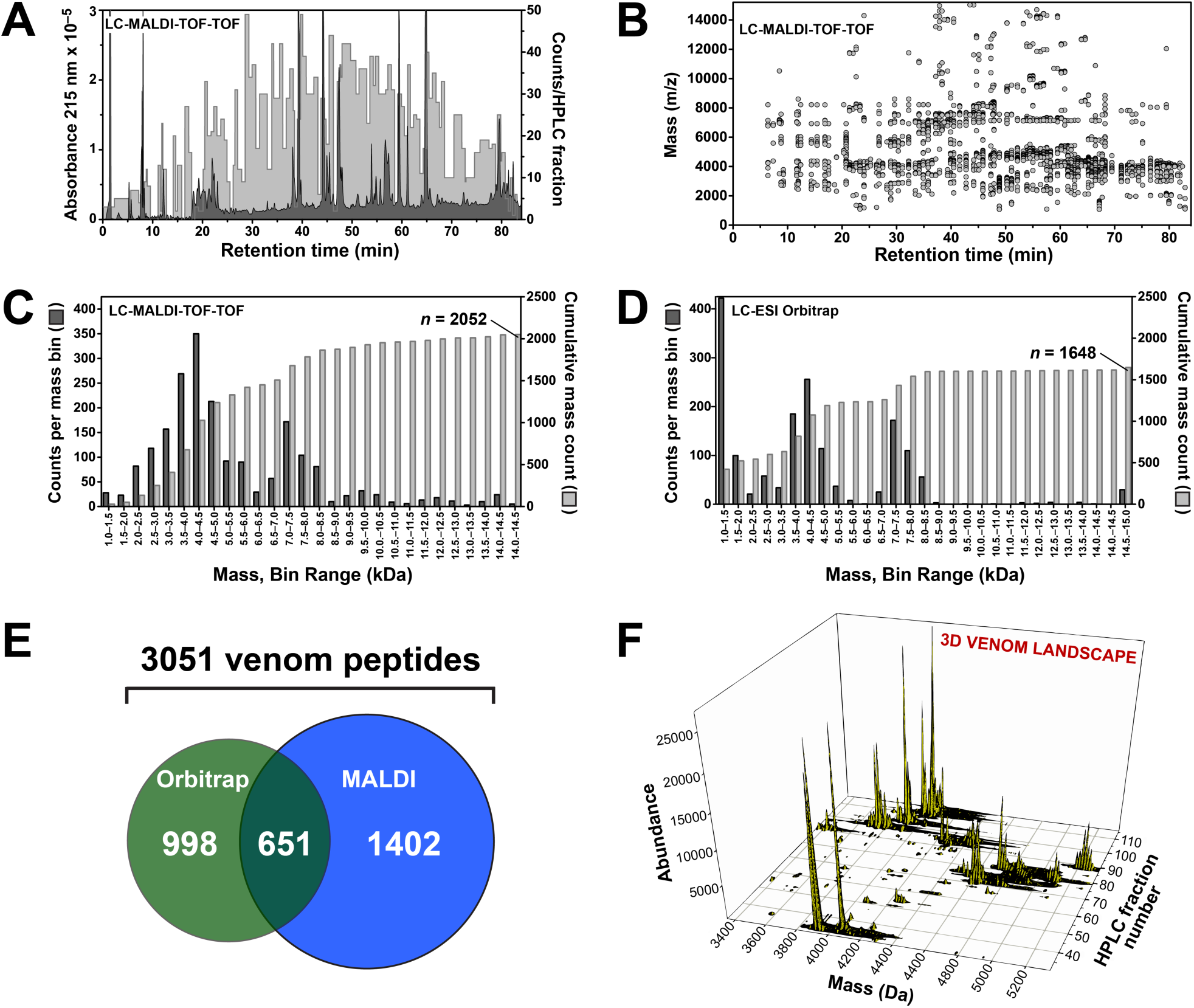
Mass profile of *H. infensa* venom. (A) RP-HPLC chromatogram showing fractionation of crude *H. infensa* venom. Absorbance at 215 nm is shown in dark grey (left ordinate-axis) while the mass count for each HPLC fraction is shown in light grey (right ordinate-axis). (B) Distribution of MALDI-TOF/TOF masses as a function of RP-HPLC retention time. (C-D) Histograms showing distribution and abundance of peptides in *H. infensa* venom as detected using (C) MALDI-TOF/TOF and (D) Orbitrap MS. Masses are grouped in 500-Da bins. Grey bars indicate the cumulative total number of toxins (right ordinate axis). (E) Euler plot showing degree of overlap between mass counts generated via MALDI-TOF/TOF and Orbitrap MS. (F) 3D landscape of *H. infensa* venom showing the correlation between RP-HPLC retention time and peptide mass and abundance generated by MALDI-TOF/TOF analysis.

The distribution of peptide masses in *H. infensa* venom is bimodal. Most peptides fall in the mass range 3.0–5.5 kDa, but there is a significant cohort of larger peptides with masses 6.5–8.5 kDa (Fig. 1C, D). This bimodal distribution matches that previously described for venom from related Australian funnel-web spiders^6,27^, various tarantulas^28^, and the spitting spider *Scytodes thoracica*^29^, and it is also reflected in the mass profile generated for all spider toxins reported to date^5^. As reported previously for Australian funnel-web spiders^6,27^, the 3D venom landscape (Fig. 1G) revealed no correlation between peptide mass and peptide hydrophobicity, as judged by reversed-phase HPLC retention time.

### Transcriptomics uncovers the biochemical diversity of the H. infensa venome

Consistent with the proteomic analysis of crude venom, sequencing of a venom-gland transcriptome from *H. infensa* revealed one of the most biochemically diverse venoms described to date, with at least 33 toxin superfamilies (Fig. 2). In light of their toxic function, each superfamily of toxins was named as suggested previously^30^ after gods or deities of death, destruction, or the underworld. Expressed sequence tags were sequenced using the 454 platform and assembled using MIRA software v3.2^31^. This produced a total of 26980 contigs and 7194 singlets, with an average contig length of 496 bp (maximum length 3159 bp, N50 674 bp). Array-based pyrosequencing (454) was used in preference to short-read sequencing to avoid ambiguous bioinformatic re-construction of highly similar toxin isoforms: the average read length was long enough to encompass entire toxin precursors, thereby avoiding sequence ambiguities. After assembly, all contigs and singlets were submitted to Blast2GO^32^ to acquire BLAST hits and functional annotations. Subsequently, sequences were grouped into three main categories: (1) proteins and other enzymes; (2) toxins and toxin-like peptides; and (3) sequences with no hits in ArachnoServer, a manually curated database of spider toxins^33^, or NCBI. 61% of all contigs/singlets were proteins/enzymes, 14% were toxins/toxin-like peptides, and 25% had no database hits (Supplementary Fig. 1S). BLAST hits included other spider toxins, as well as peptides and proteins found in other venomous arthropods with recently annotated genomes, including the tick *Ixodes scapularis,* the honeybee *Apis mellifera,* the jumping ant *Harpegnathos saltator*, and the parasitoid wasp *Nasonia vitripennis*.

**Figure 2:**
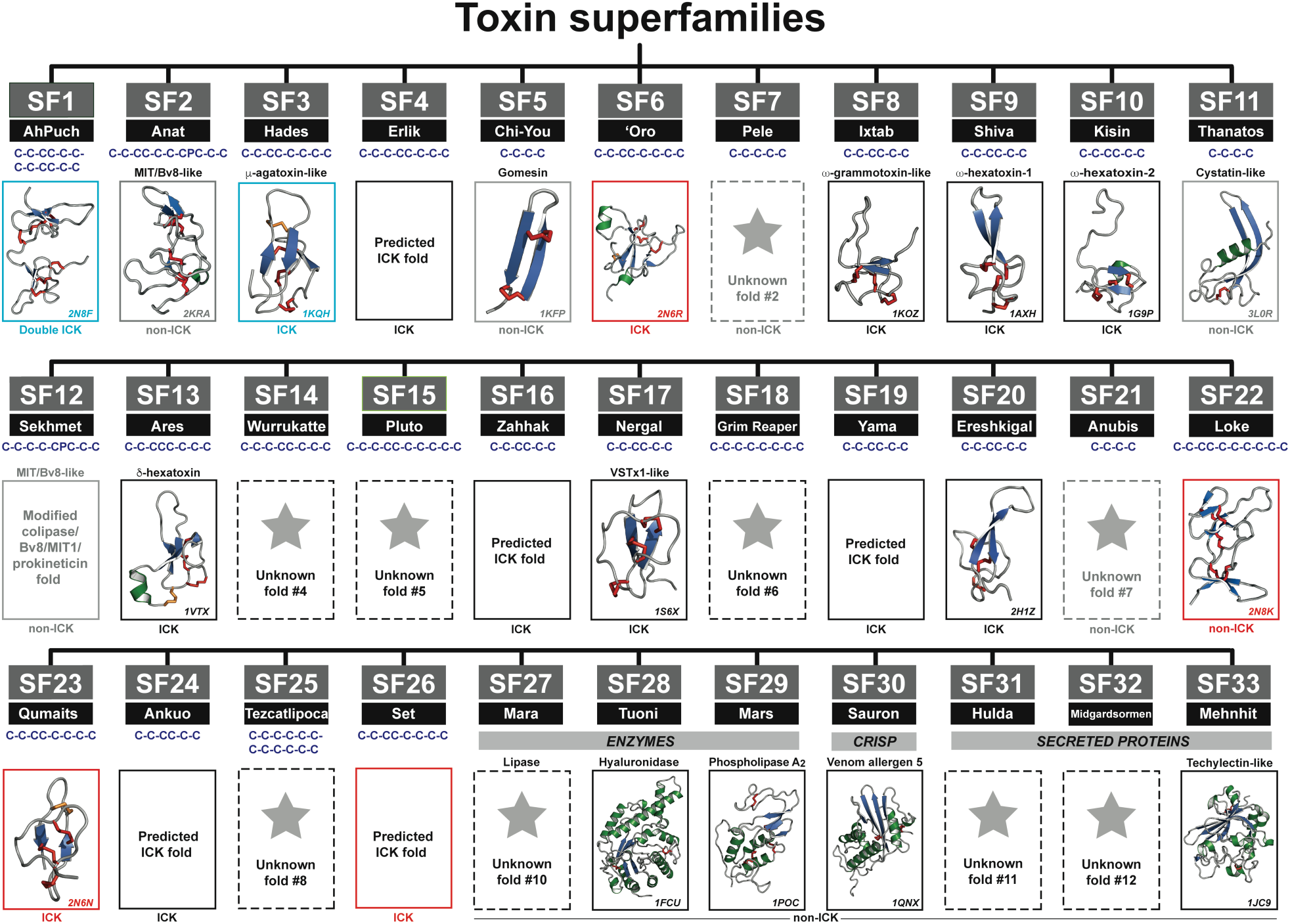
Overview of the venom proteome of *H. infensa*. The venom proteome of *H. infensa* consists of 33 toxin superfamilies (SF1–SF33). For each SF, the cysteine framework is shown in blue, and the 3D fold is classified as ICK, putative ICK, double ICK, non-ICK, or unknown. Light blue boxes enclose previously solved structures of superfamily members from *H. infensa*. Black boxes enclose structures of orthologous superfamily members from related mygalomorph spiders or, in the case of enzymes and CRiSP proteins, orthologs from venomous hymenopterans (bees and wasps). Red boxes enclose structures solved in the current study (SF6, SF22, SF23, and SF26). Stars enclosed by dashed black boxes signify toxin superfamilies for which no structural information is currently available. For each of the structures, β strands are shown in blue, helices are green, and core disulfide bonds are shown as solid red tubes. For DRPs containing an ICK motif, additional disulfide bonds that do not form part of the core ICK motif are shown as orange tubes. Toxin superfamilies are named after gods or deities of death, destruction and the underworld. For each structure shown, Protein Data Bank accession numbers are given in the lower right corner.

To maximise the discovery of novel DRPs, an in-house algorithm was written to allow discovery of potential open reading frames (ORFs). Both annotated and novel peptides were then combined into a single file that was used to analyse the processing signals in all precursors. First, for each toxin group, signal peptide cleavage sites were predicted using SignalP v4.0^34^ while propeptide cleavage sites were predicted based on a sequence logo analysis^35^ of all known spider-toxin precursors documented on ArachnoServer (Supplementary Fig. 2S). After determination of signal and propeptide regions, toxins were classified into superfamilies (SF) based on their signal-peptide sequence, total number of cysteine residues, and their known or predicted cysteine framework, as shown in Figs 2 and 3.

**Figure 3:**
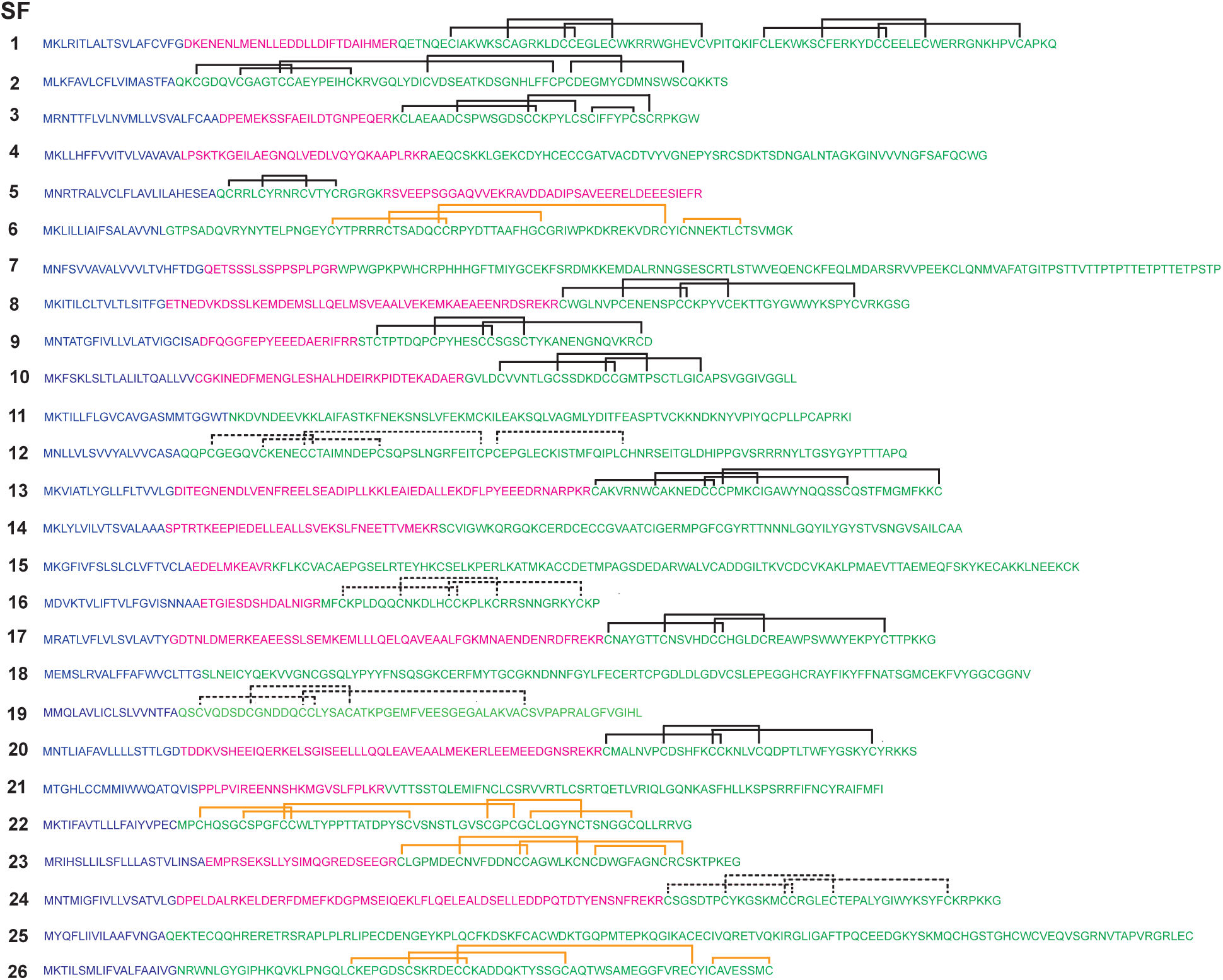
DRP superfamilies in *H. infensa*. Consensus sequences for each of the toxin superfamilies in *H. infensa* venom. Signal peptide, propeptide, and mature toxin sequences are shown in blue, magenta, and green, respectively. The lines above each sequence indicate the disulfide framework, with solid and dashed lines indicating disulfide bonds that were determined experimentally (including via the structure of close homologues) or predicted, respectively. Orange lines correspond to disulfide frameworks that were determined experimentally in the current study.

Transcript abundances for each superfamily were estimated using RSEM v1.2.31^36^. Briefly, trimmed reads were mapped back to the assembled transcriptome. We grouped all the identifiers of contigs containing sequences that encode toxins/proteins from each superfamily; the list of identifiers was saved on a text file and used to search the RSEM output. Transcripts Per Million (TPM), were used to estimate the transcript abundance for each superfamily and the total number of transcripts obtained are summarized in Table 1S and Figure 4. Differences in transcript abundances vary from low TPM counts for superfamilies 11, 21 and 28 to the most abundantly expressed peptide in the venom gland, SF4, which interestingly had not been previously characterised as a major component of the venom of *H. infensa*.

**Figure 4:**
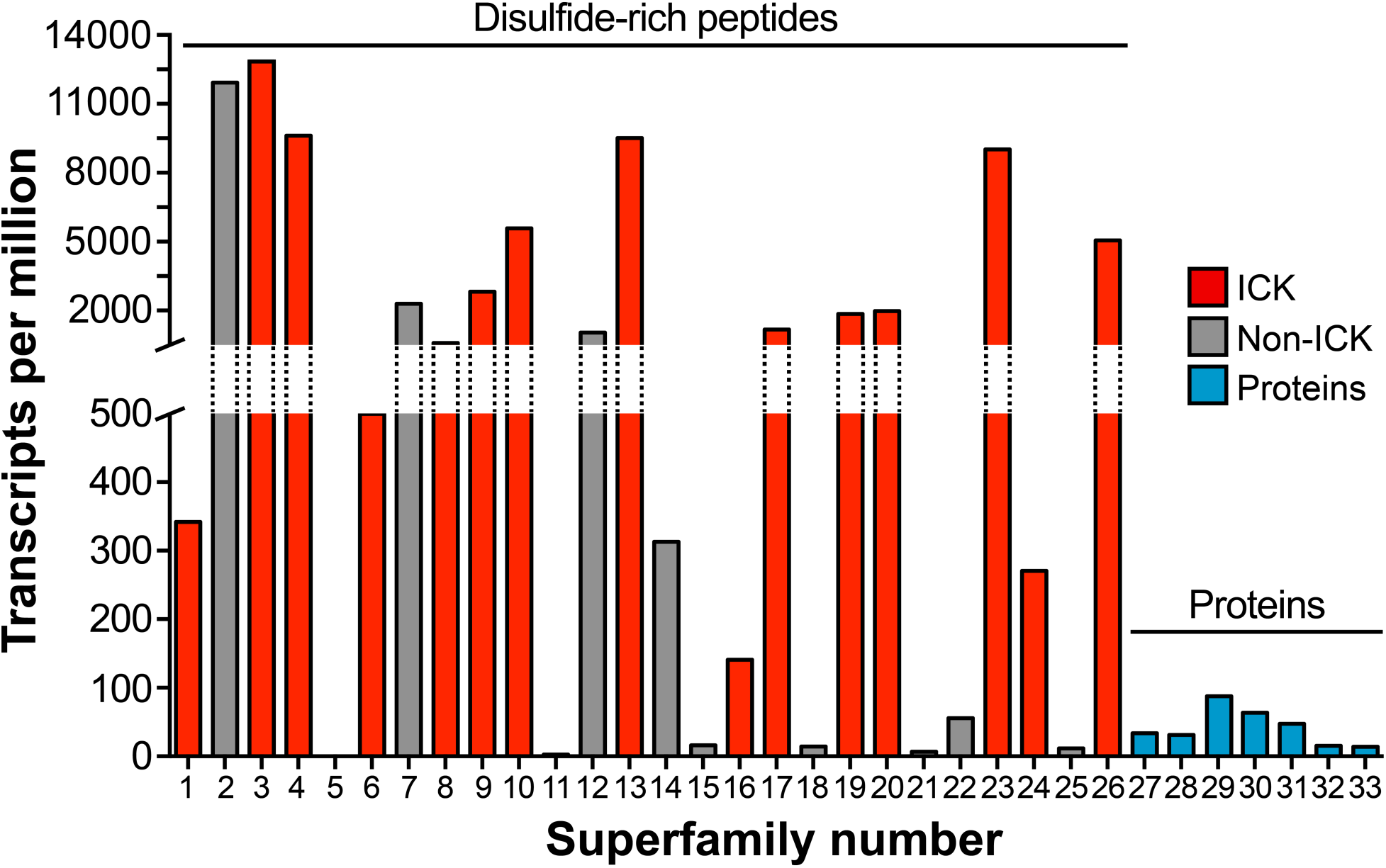
Mass profile of *H. infensa* venom. Abundance of transcripts encoding DRPs and proteins obtained from sequencing of a *H. infensa* venom-gland transcriptome. Blue bars represent protein-encoding transcripts, while red and grey bars denote transcripts encoding DRPs that do or don’t have an ICK scaffold, respectively. ICK transcripts dominate the venom-gland transcriptome. Superfamily numbers correspond to those shown in Figure 2.

Using this approach, we determined that the venom gland of *H. infensa* expresses at least 26 families of DRPs, three families of enzymes (hyaluronidase, lipase, and phospholipase A_2_ (PLA_2_)), one family of cysteine-rich secretory proteins (CRiSPs), and three families of secreted proteins (Figs 2 and 3). A brief description of each superfamily is provided in the supplementary material. Hyaluronidase and PLA_2_ have both been convergently recruited into many animal venoms^37^. Venom hyaluronidases act as diffusion factors that enhance tissue permeability to facilitate spreading of neurotoxins^38^ or hemostatic factors^37^. Sequence diversity across taxa is minimal and no new activities have been reported for venom hyaluronidases^37^. In contrast, venom PLA_2_s are more diverse and they have acquired a range of new functions, some of which are independent of their catalytic activity^39,40^. There are only a few reports of lipases in animal venoms, but the *H. infensa* venom lipase has significant homology to those found to be active in wasp venoms^41,42^. It is unclear what function lipases might play in spider venom.

### Proteomic analysis of milked venom confirms the biochemical complexity of H. infensa venom

To further probe the proteomic complexity of the venom of *H. infensa*, we searched our annotated venom gland transcriptome with tandem mass spectrometry spectra generated from fractionated, reduced, alkylated and trypsin digested venom. Using a stringent confidence threshold corresponding to a < 1% false discovery rate calculated from a decoy-based false discovery rate analysis in ProteinPilot, we were able to detect 1109 of the 1225 venom proteins and peptides predicted from our transcriptomic analyses.

These proteomically identified venom components spanned 22 of the predicted 33 superfamilies, including all of the most diverse superfamilies, and three of the 7 predicted protein superfamilies. There was also a strong correlation between low expression and lack of proteomic detection; the predicted peptide superfamilies with a cumulative expression value less than 100 TPM accounted for all of the peptide superfamilies not detected in the venom (Fig. 4). Taken together, our mass spectrometry, transcriptomic, and proteomic analyses confirm that the venom of *H. infensa* is a peptide-dominated venom of exceptional biochemical complexity.

### Actively regenerating spider venom glands are enriched in toxin processing machinery

We depleted the venoms glands of spiders by electrical stimulation three days before sacrificing them to obtain venom glands for transcriptomic analysis. These venom glands were therefore actively engaged in venom regeneration at the time of sampling. Thus, as expected, the venom-gland transcriptome was highly enriched in transcripts encoding proteins involved in translation, processing, folding and posttranslational modification of venom toxins.

Non-toxin contigs/singlets were classified into EuKaryotic Orthologous Groups (KOGs) and classified according to gene ontology (GO). Blast2GO was used to generate summaries for the most common GO types: ‘biological’, ‘cellular’ and ‘molecular’ processes (Supplementary Figure 3S). The most abundant GO terms within the ‘cellular process’ category were protein-binding (i.e., zinc finger proteins) and the cytoskeleton. Similarly, regulators of high-energy phosphatases, cytochrome oxidases, and NADH dehydrogenase (i.e., components of the electron transport chain) were also found, consistent with a biochemically active venom gland.

Within the ‘biological process’ classification, metabolic and cellular processes accounted for 28% and 32% of the GO terms, respectively. Within these classes we found transcripts encoding translocases, cyclases, elongation factors, convertases, histones, reverse transcriptases, aminotransferases, tRNA synthetases, translation initiation-factors, ribosomal proteins, polyadenylate binding proteins, helicases, RNA polymerase, and rRNA promoter binding proteins. These proteins are involved in transcription and translation of peptides, processes expected to be highly active in a venom gland producing a large number of venom-related transcripts.

Finally, within the ‘molecular process’ classification, binding and catalytic activity were the most abundant GO terms listed. A large number of identified contigs/singlets included in these classifications encode enzymes related to protein folding, transport, ubiquitination, post-translational modification, and proteolysis. The list included protein disulfide isomerase, carboxypeptidases, chaperonins, heat shock proteins, signal peptide peptidases, gluthathione *S*-transferases, dismutases, and thioredoxins, all of which are important for the processing and maturation of venom proteins and DRPs.

### The venom of *H. infensa* contains both classical and highly elaborated ICK toxins

Within the venom-gland transcriptome of *H. infensa*, 26 of the 33 toxin superfamilies were identified as DRPs (Fig. 2). Thus, DRPs are the dominant proteins in Australian funnel-web spider venom, even when transcript abundance is taken into account (see below and Fig. 4). Seven of the 26 DRP superfamilies (SFs) could be confidently classified as knottins based on prior determination of the 3D structure of one of the superfamily members (SF3) or strong homology with an orthologous knottin (i.e., from another spider venom) whose structure has been elucidated (SFs 8–10, 13, 17, and 20). A further four superfamilies were confidently predicted to be knottins because their cysteine framework and inter-cysteine spacing conform to the ICK motif (SFs 4, 16, 19 and 24), albeit with an additional disulfide bond in the case of SF4. In addition, SF1 is a family of double-knot toxins, one of whose structure was recently solved (Fig. 5A)^22^. Thus, based on prior knowledge, 12 of the 26 DRP superfamilies can be assigned as knottins. Only six DRP superfamilies could be confidently assigned as not being knottins: SF2 corresponds to the MIT/Bv8/prokineticin superfamily^43,44^, SF5 is a family of β-hairpin toxins with high homology to the antimicrobial peptide gomesin isolated from hemocytes of the tarantula *Acanthoscurria gomesiana*^45^, SF11 is a family of cystatin-like toxins, SF12 is a derived MIT/Bv8/prokineticin family with four rather than the canonical five disulfide bonds, and SF7 and SF21 have less than the minimum of six cysteine residues required to form an ICK motif.

**Figure 5:**
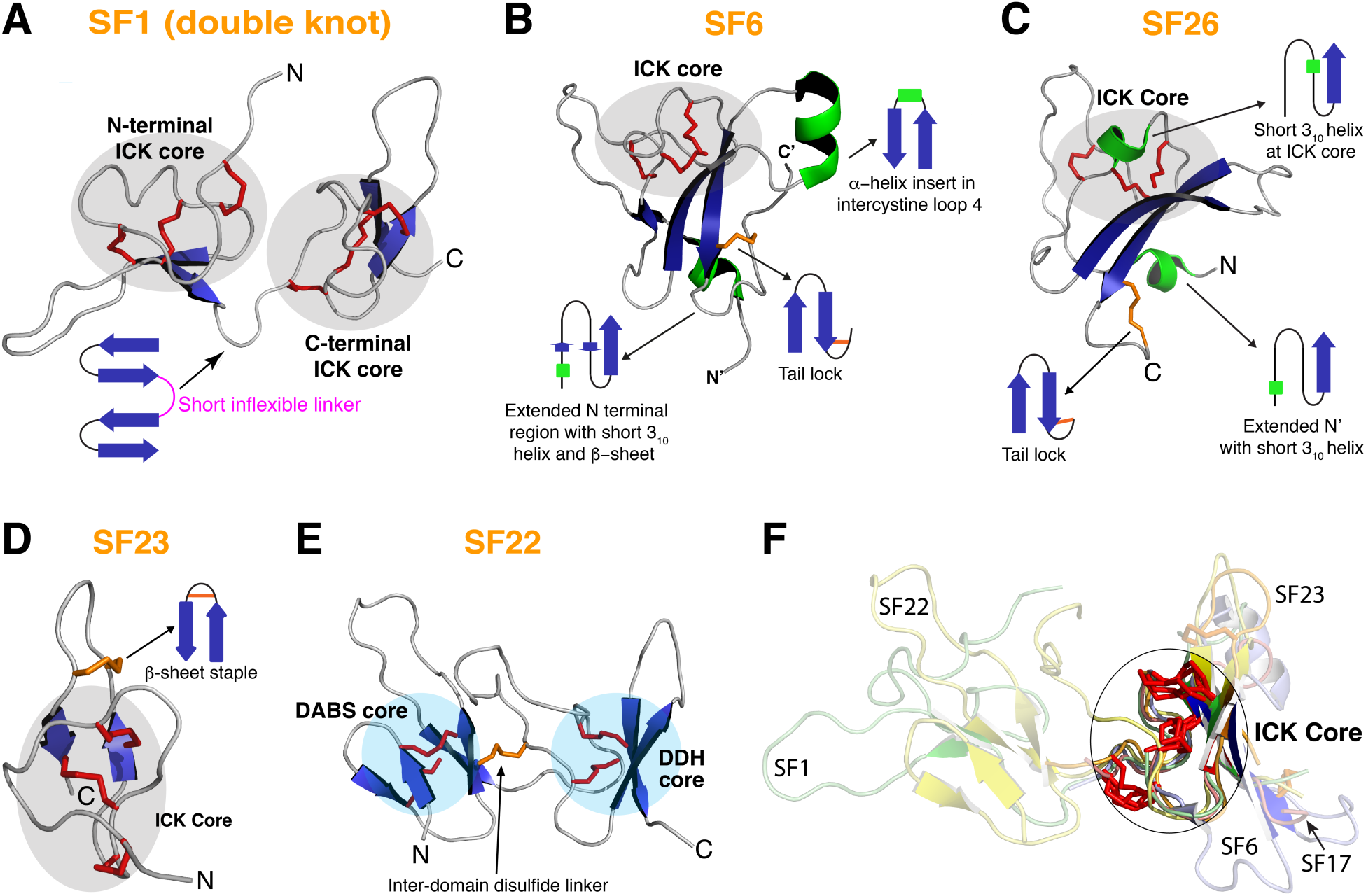
3D structures of selected DRPs found in the venom of *H. infensa*, highlighting some of the key structural innovations in “short” (<5 kDa) and “long” venom peptides (6-10 kDa) (A) Schematic representation of the SF1 double-knot toxin which is comprised of two independently folded ICK domains joined by an inflexible linker. (B) SF6 family peptides adopt a typical ICK fold with several unique elaborations, including an extended C-terminal tail that is stapled to the rest of the structure via an additional disulfide bond (“tail-lock”), an enlarged intercystine loop 4 that contains an α-helical insertion, and a highly extended N-terminal region that includes a 6-resiude α-helix and a short two-stranded β-sheet. (C) SF26 DRPs also adopt a highly elaborated knottin fold with a C-terminal tail tail-lock, a 5-residue 3_10_ helix in the core ICK region between Cys^II^ and Cys^III^, and a long N-terminal extension that includes a short α-helix. (D) SF23 DRPs only differ from classical ICK toxins by having an extra disulfide bond (“β-sheet staple”) that stabilizes the β-hairpin loop. (F) SF22 DRPs represents an entirely new toxin fold comprised of two independent structural domains connected by a disulfide bond. The C-terminal domain forms a DDH core while the N-terminal domain adopts a newly described <u>d</u>isulfide-stabilized <u>a</u>ntiparallel β-hairpin <u>s</u>tack (DABS). In panels A–E, disulfide bonds comprising the ICK motif are shown as red tubes, while additional non-core disulfide bonds are shown as orange tubes. β sheets and α-helices are highlighted in blue and green, respectively. (E) Structural alignment of the ICK core regions of DRPs from SFs 1, 6, 17, 22 and 23. highlighting the strong conservation of the ICK motif (DDH in the case of SF22) regardless of the extent of structural elaborations outside this core region.

Eight of the 26 DRP superfamilies with three or more disulfide bonds have novel sequences with unknown structure and biological activity (SFs 6, 14, 15, 18, 22, 23, 25, and 26). Thus, in order to further explore the structural diversity and evolutionary origin of spider-venom DRPs, we attempted to determine the structure of representative members of SFs 6, 14, and 26. We also produced homologous sequences to those of SFs 22 and 23. These superfamilies were prioritised over SFs 15, 18, and 25 which contain only low-abundance transcripts (Fig. 4), indicating they are minor components of the venom. Peptides were expressed in the periplasm of *E. coli* using a system we previously optimized for production of DRPs^46,47^, then purified using nickel-affinity chromatography followed by reversed-phase HPLC (Supplementary Fig. 4S). All of the chosen peptides were successfully produced with the exception of the representative member of SF14, which failed to express in *E. coli* for reasons that remain to be determined. The SF6, SF22, SF23, and SF26 peptides were uniformly labelled with ^15^N and ^13^C, and their structures determined using a 3D/4D NMR strategy we described previously that takes advantage of non-uniform sampling to expedite data acquisition and improve structure quality^48^. Each of the structures we determined are of high precision and have excellent stereochemical quality as evidenced by the structural statistics presented in Supplementary Table S2.

The SF6 family member adopts a highly derived knottin fold with several unique elaborations (Fig. 5B): (i) the β-hairpin loop encompasses an 8-residue α-helix; (ii) an extended C-terminal tail is locked in place by a disulfide bridge to the adjacent C-terminal β-strand of the ICK motif; and (iii) a highly extended N-terminal region (i.e., preceding the first cysteine residue) includes a short 6-residue α-helix and a β-strand that forms a small β-sheet via hydrogen bonds with residues in the Cys^IV^-Cys^V^ intercystine loop. The SF26 family member also adopts a highly modified knottin fold (Fig. 5C), with the following elaborations of the classical ICK motif: (i) similar to SF6 toxins, an extended C-terminal tail is stapled in place by a disulfide bridge between the C-terminal Cys residue and a Cys residue at the base of the adjacent C-terminal β-strand of the ICK motif; (ii) the long N-terminal region includes a short 6-residue 3_10_ helix followed by an extended segment that stretches across towards the much longer β-sheet of the cystine knot; and (iii) a 5-residue 3_10_ helix is inserted in the Cys^II^–Cys^III^ loop in the ICK motif. The SF23 family member adopts a prototypical knottin fold with an additional stabilizing disulfide bond at the base of the β-hairpin loop (Fig. 5D), which is a relatively common elaboration in spider toxins^49^. The new structures of representative toxins from superfamilies SF6, SF23, and SF26 highlight the fact that spiders have evolved many different elaborations of the ancestral ICK fold in order to diversify their venom arsenal.

Elucidation of the 3D structure of a toxin (U_33_-TRTX-Cg1c) from the Chinese tarantula *Chilobrachys guanxiensis* which is homologous to SF22 revealed that these toxins do not adopt an ICK fold. The overall topology of SF22 (Fig. 5E) is similar to the prokineticin/colipase fold adopted by other five-disulfide DRPs such as the SF2 and SF12 toxins, but they are not homologous. The prokineticin/colipase fold found in vertebrate proteins can be viewed as a head-to-tail duplication of two DDH motifs with various elaborations on the loops^10^. SF22 also contains two core motifs. The C-terminal motif is structurally similar to the C-terminal subdomain of prokineticin and colipase, and it adopts a DDH motif with a long 17-residue loop between Cys^VI^ and Cys^VIII^ (c.f. 4–6 residues in the equivalent loop of most ICK motifs). The extended loop bridges over to the N-terminal motif, stabilised by an inter-subdomain disulfide bond (Cys^III^-Cys^VII^) and a hydrogen bond between Cys15 (Cys^IV^) and Tyr49. However, in contrast with prokineticin and colipase, the N-terminal region does not adopt a DDH fold. Rather, the N-terminal core is composed of two stacked antiparallel β-hairpins in which the parallel strands in the adjacent hairpins are covalently linked by two disulfide bonds (Fig. 5E). We have named this N-terminal core motif of SF22 the <u>D</u>isulfide-stabilized <u>a</u>ntiparallel β-hairpin <u>s</u>tack, DABS. This peculiar motif is unprecedented in venom-peptides, making this DABS-DDH motif a novel class of toxin.

Superimposition of the 3D structures of SF6, SF23, SF26 and either one of the two ICK domains of SF1 reveals that, regardless of the size and structural elaborations acquired in these peptides (Fig. 5F), the central ICK motif remains essentially the same, despite enormous variations in amino acid sequence.

### ICK toxins dominate the venom arsenal of H. infensa

Our combined transcriptomic, proteomic, and structural biology data revealed that ICK toxins are the major contributor to the diversity of the *H. infensa* venom peptidome, with 15 of the 26 identified DRP superfamilies comprised of simple or highly derived knottins. Moreover, analysis of transcript abundance revealed that 11 of the 14 most highly expressed DRP superfamilies are ICK toxins (Fig. 4). Overall, based on transcript abundance, ICK toxins represent ~91% of the total venom peptidome. If one includes the seven superfamilies of putative protein toxins (SF27–SF33), which are all expressed at very low levels (Fig. 4), the venom proteome is composed of 90.5% ICK toxins, 9.1% non-ICK DRPs, and 0.4% protein toxins. We conclude that ICK toxins dominate the venom arsenal of *H. infensa* both in terms of molecular diversity and abundance.

### Phylogenetic analyses suggests a single early recruitment of ICK toxins in H. infensa

In order to investigate whether the diverse and abundant ICK toxins in spider venom have a common origin, we examined their phylogenetic relationship with homologous sequences from non-venom gland spider tissues (i.e., hypodermal tissue from *Cupiennius saliei* and leg tissue from *Liphistius malayanus* and *Neoscona arabesca*) and tissue from a non-venomous uropygid arachnid (*Mastigoproctus giganteus*) (Fig. 6). BLAST searches of all identified spider venom ICK variants against these non-venom datasets returned only hits to venom peptides with classical ICK folds, suggesting the elaborate ICK structures observed for several of the venom peptide superfamilies found in *H. infensa* represent extreme cases of structural diversification following functional recruitment as toxins. Although we were unable to resolve the higher-order phylogenetic relationships between the ICK toxin superfamilies, both our Bayesian inference and maximum likelihood reconstruction analyses revealed that the venom knottins form a well-supported monophyletic clade (Fig. 6). This suggests that the structural diversity of ICK toxins, which make up the bulk of the molecular diversity in the venom of *H. infensa*, probably arose after the functional recruitment of a “classical” ICK fold into the venom of an ancestral spider.

**Figure 6:**
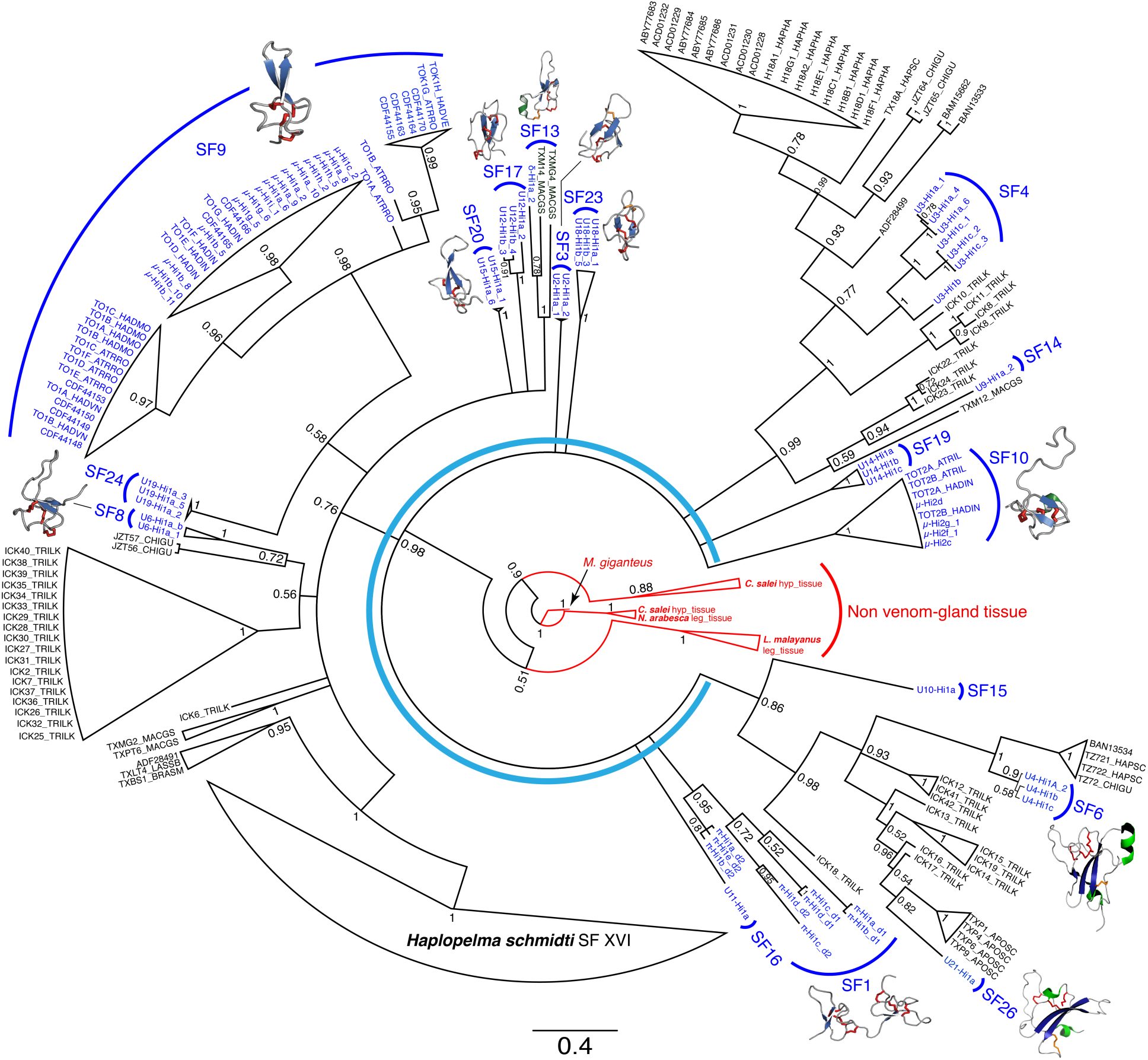
Evolution of spider-venom ICK toxins. Maximum likelihood tree showing the phylogenetic relationship between *H. infensa* DRPs and DRPs isolated from the venom gland or other tissues of other spider species. The tree was rooted using the whip scorpion *Mastigoproctus giganteus* as the outgroup. Bootstrap values are shown at each node. The tree shows that although many of the phylogenetic relationships between superfamilies remains unresolved all venom-derived DRPs form a well-supported monophyletic clade. Superfamily sequences belonging to *H. infensa* are highlighted in blue text and representative structures for each superfamily are shown. DRP sequences from muscle and other tissues are highlighted in red; all other sequences (denoted by the light blue broken circle) represent venom DRPs Venom peptides isolated from other species have their corresponding accession numbers/common toxin names listed in the labels with exception of the *H. schmidti* superfamily XVI clade. A summary of all accessions used can be found in the supplementary material.

Although the phylogenetic relationships between many of the ICK superfamilies remain unresolved, our analyses provide insight into the loss and gain of disulfide bonds in these toxins (Fig. 6). Although the core ICK motif contains only three disulfide bonds^12^, our data suggest that the ancestral ICK spider toxin either contained a fourth disulfide bond stabilising the β-sheet in loop 4, or that this additional disulfide was gained on at least two separate occasions. A loss of intramolecular disulfide bonds may seem counterintuitive during the evolution of venom DRPs for which stability is paramount. However, intriguingly, the stabilising loop-4 disulfide bridge was present in all homologues identified in the outgroup datasets (data not shown). Moreover, disulfide loss appears to have been crucial during evolution of DDH toxins^11^ from ICK precursors in scorpions^23^, suggesting that this scenario may not be as unlikely as previously postulated^11,50^.

Regardless of whether the ancestral ICK spider toxin contained the loop-4 disulfide, our data indicate that *gain* of disulfide bonds underlies the myriad of structural variations in ICK spider toxins. This not only highlights the importance of disulfide bonds in stabilising the structure of spider toxins but further illustrates the extraordinary versatility of the ICK fold. The permissiveness of the ICK fold to both sequence variation^23^ and disulfide elaborations likely explains why it appears to be the most widespread cysteine-rich fold known, being found in a diverse range of taxa including arachnids, fungi, insects, molluscs, plants, sea anemones, sponges, and even viruses^23,51^. ICK peptides also appear to constitute the most abundant fold in the “dark proteome”^52^, suggesting that we still have much to learn about their structural and functional plasticity.

### Conclusions

Our holistic approach to the analysis of *H. infensa* venom has revealed the likely pathway for molecular evolution of venom in Australian funnel-web spiders and mygalomorph spider more generally. Multiple duplications of a single ancestral ICK gene were followed by periods of diversification, the generation of structural variants, and the insertion of post-translational modifications, with selection of variants via adaptive evolution to generate the bulk of small venom DRPs. There appears to have limited intragenic gene duplication to produce double-knot toxins such as the SF1 family, with most venom peptides with higher masses (from 6–10 kDa) having arisen from elaborations of a single ICK scaffold, often accompanied by the inclusion of additional disulfide bonds (Fig.7). In summary, our work suggests that the extraordinary pharmacological complexity of spider venoms is largely due to a panel of structurally related DRPs that all evolved from a single ancestral ICK toxin.

**Figure 7:**
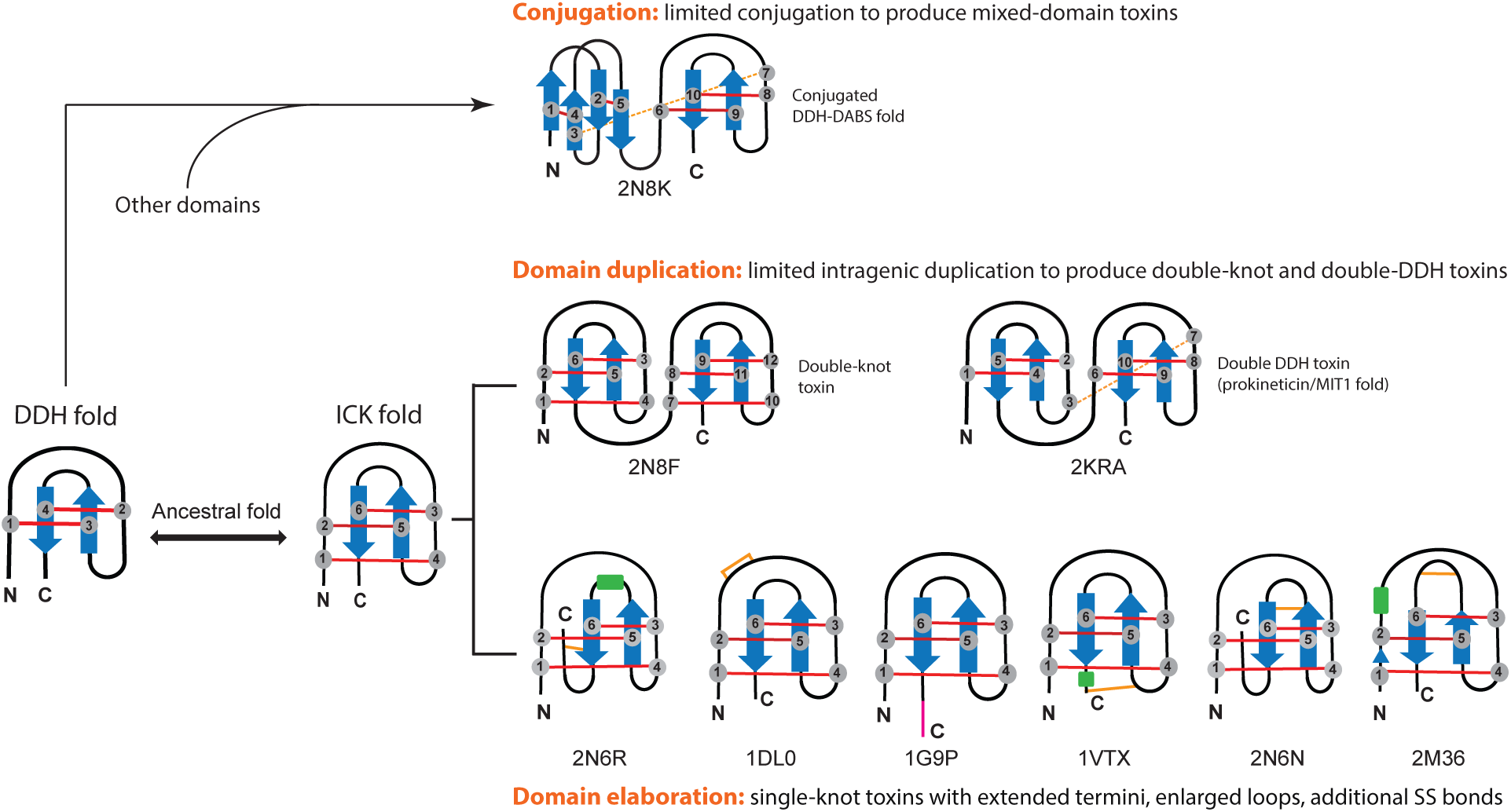
Overview of structural innovations in spider-venom ICK peptides. Schematic overview of the mechanisms by which an ancestral ICK or DDH toxin was duplicated, conjugated and elaborated upon to form the diversity of ICK scaffolds found in extant mygalomorph spider venoms. Gray numbered circles represent cysteine residues, with core disulfide bonds that form the cystine knot or DDH motif indicated by solid red lines. Additional non-core disulfides are highlighted in orange, with dashed lines indicating inter-domain disulfide bonds. Blue arrows and green rectangles denote β-strands and α-helices, respectively. N- and C-termini are labelled, and Protein Data Bank accession codes for representative structures are given below each schematic peptide fold.

## Materials and Methods

### Spider collection

Specimens of *Hadronyche infensa* (Hickman) were collected from Orchid Beach, Fraser Island, Queensland, Australia. Spiders were housed individually at 23–25°C in plastic containers (approximately 155 × 155 × 140 mm) in dark cabinets.

### Venom collection

Spiders were aggravated using a pair of forceps until venom was expressed on the fang tips. Venom was then collected by aspiration and stored at –20°C until needed. If impurities such as soil or sand were detected, the venom was diluted to 10 times its initial volume with ultrapure water and separated from contaminants using a low-protein-binding 0.22 µm Ultrafree-MC centrifugal filter (Millipore, MA, USA).

### Venom profiling

The venom profile of the Australian funnel-web spider *H. infensa* was obtained using a combination of reversed-phase (RP) HPLC and MS techniques^6^. Offline LC-MALDI-TOF/TOF was utilized in order to maximize the amount of mass information extracted. A conservative analysis was used to manually curate MS datasets. Analysis criteria included: (a) only spectra with well-defined peaks were considered as “real”; (b) a signal-to-noise ratio of 15 was set before any masses were recorded; (c) masses were excluded if they met the following criteria relative to another recorded mass: +16 Da and +32 Da (oxidized), –18 Da (dehydrated), –17 Da (deamidated), +22 Da (sodium adduct); (d) masses <1000 Da, which were presumed to correspond to small organic compounds such as polyamines or matrix clusters, rather than peptides, were excluded from the analysis. Duplicated masses were eliminated from adjacent fractions and a final list of sorted masses was generated. Potential dimers and doubly charged species were also eliminated (± 3–5 Da). 3D contour plots, or ‘venom landscapes’, were constructed from LC-MALDI-TOF/TOF MS data as described previously^6^.

For proteomic analysis of venom composition, 5 mg of milked venom pooled from three spider specimens was fractionated on a Shimadzu Prominence HPLC (Kyoto, Japan) using a Vydac C_18_ RP-HPLC column (300 Å, 5 µm, 4.6 mm × 250 mm). Venom was eluted across a gradient of 5–80% solvent B (90% acetonitrile [ACN], 0.043% trifluoroacetic acid [TFA]) in solvent A (0.05% TFA) over 120 min. Twelve fractions were manually collected (one every 10 min) and absorbance was monitored at 214 nm and 280 nm. Fractions were lyophilized, reconstituted in ultrapure water, and 1/10 removed for proteomic analyses. Fractionated proteins and peptides were reduced with dithiotheitol (5 mM in 50 mM ammonium bicarbonate, 15% ACN, pH 8), alkylated with iodoacetamide (10 mM in 50 mM ammonium bicarbonate,, 15% ACN, pH 8), and digested by overnight incubation with trypsin (30 ng/µL in 15 µL 50 mM ammonium bicarbonate, 15% ACN, pH 8). Upon completion of digestion, formic acid (FA) was added to a final concentration of 5% and the digested samples desalted using a C18 ZipTip (Thermo Fisher, Waltham, MA, USA).

Desalted, digested samples were dried in a vacuum centrifuge, dissolved in 0.5% FA, and 2 µg of each analysed by LC-MS/MS on an AB Sciex 5600TripleTOF (AB Sciex, Framingham, MA, USA) equipped with a Turbo-V source heated to 550°C and coupled to a Shimadzu Nexera UHPLC. Samples were fractionated on an Agilent Zorbax stable-bond C18 column (2.1 × 100 mm, 1.8 µm particle size, 300 Å pore size), across a gradient of 1–40% solvent B (90% ACN, 0.1% FA) in 0.1% FA over 45 min, using a flow rate of 180 µL/min. MS1 survey scans were acquired at 300–1800 *m/z* over 250 ms, and the 20 most intense ions with a charge of +2 to +5 and an intensity of at least 120 counts/s were selected for MS2. The unit mass precursor ion inclusion window was ± 0.7 Da, and isotopes within ± 2 Da were excluded from MS2, which scans were acquired at 80–1400 *m/z* over 100 ms and optimized for high resolution. For protein identification, MS/MS spectra were searched against the translated annotated venom gland transcriptome of *H. infensa* using ProteinPilot v4.0 (AB Sciex, Framingham, MA, USA). Searches were run as thorough identification searches, specifying tryptic digestion and cysteine alkylation by iodoacetamide. Decoy-based false discovery rates (FDR) were estimated by ProteinPilot, and for our protein identification we used a protein confidence cut-off corresponding to a local FDR of < 1%. Spectra were also manually examined to further eliminate any false positives.

### Library construction and sequencing

Three spiders were milked by electrical stimulation to deplete their glands of venom then, three days later, they were anesthetized and their venom glands dissected out and immediately placed in TRIzol^®^ reagent (Life Technologies). Total RNA was extracted following the standard TRIzol^®^ protocol. mRNA enrichment from total RNA was performed using an Oligotex direct mRNA mini kit (Qiagen). RNA quality and concentration was measured using a Bioanalyzer 2100 pico chip (Agilent Technologies).

A cDNA library was constructed from 100 μg mRNA using the standard Roche cDNA rapid library preparation and emPCR method. Sequencing was carried at the Australian Genome Research Facility (AGRF-Brisbane) using a ROCHE GS-FLX sequencer. The Raw Standard Flowgram File (.SFF) was processed using Cangs software, and low quality sequences discarded (Phred score cut-off of 25)^53^. *De novo* assembly was performed using MIRA software v3.2^31^ using the following parameters: -GE:not=4 -- project=Hinfensa --job=denovo,est,accurate,454 454_SETTINGS -CL:qc=no -AS:mrpc=1 - AL:mrs=99,egp=1. The assembled dataset was visualized using Geneious software (www.geneious.com). Consensus sequences from contigs and singlets were submitted to Blast2GO (www.blast2go.com) to acquire BLAST and functional annotations^32^. In parallel to the functional annotation analysis, an in-house algorithm (Toxin|seek) was written to allow prediction of potential toxin ORFs that may represent novel DRPs. Once annotated and predicted, lists were merged, redundancies were removed, and signal sequences determined using SignalP^34^. Putative propeptide cleavage sites were predicted using a sequence logo analysis^35^ of all known spider precursors (Supplementary Fig. 2S). After identifying all processing signals, toxins were classified into superfamilies based on their signal sequence and cysteine framework. Finally, toxin abundance was estimated using RSEM v1.2.31 using the following commands: rsem-prepare-reference --bowtie2 RL5_Hinfensa.fasta Hinfensa and rsem-calculate-expression --bowtie2 --fragment-length-mean 588 Hinfensa_2010_trimmed.fastq Hinfensa Hinfensa_expression.results bowtie2 -q --phred33 --sensitive --dpad 0 --gbar 99999999 --mp 1,1 --np 1 --score-min L,0,-0.1 -p 1 -k 200 -x Hinfensa -U Hinfensa_2010_trimmed.fastq.

### Nomenclature

Toxins were named using the rational nomenclature described previously^54^. Spider taxonomy was from the World spider catalog v19.5 (https://wsc.nmbe.ch).

### Peptide expression and purification

Genes encoding toxins of interest were synthesized using codons optimized for *E. coli* expression and subcloned into a pLICC_D168 vector by GeneArt (Regensburg, Germany). Plasmids were transformed into *E. coli* BL21 (λDE3), and toxin expression and purification carried out using published methods^46,47^ (Supplementary Fig. 3). Peptides were separated from salts, His_6_-MBP and His_6_-TEV protease using a Shimadzu Prominence HPLC system (Kyoto, Japan). Separation was performed on a Jupiter C_4_ reverse phase HPLC column (250 × 10 mm, 300 Å, 10 µm; Phenomenex), using a flow rate of 5 mL/min and an elution gradient of 5–60% solvent B over 30 min. Absorbance was monitored at 214 nm and 280 nm and fractions were manually collected. Purified peptides were lyophilized, reconstituted in water or buffer, and approximate peptide concentration determined from absorbance at 280 nm measured using a NanoDrop spectrophotometer (Thermo Scientific). Electrospray ionization (ESI)-MS spectra were acquired using an API 2000 LC/MS/MS triple quadrupole mass spectrometer. Mass spectra were obtained using positive ion mode over *m*/*z* range of 400–1900 Da.

### Structure determination

The structure of toxins in superfamily SF6, SF22 SF23 and SF26 was determined using heteronuclear NMR spectroscopy. Each sample contained 300 µL of ^13^C/^15^N-labelled peptide (100–300 µM) in 20 mM 2-(*N*-morpholino)-ethanesulfonic acid (MES) buffer, 0.02% NaN_3_, 5% D_2_O, pH 6. NMR data were acquired at 25^°^C on an Avance II+ 900 MHz spectrometer (Bruker BioSpin, Germany) equipped with a cryogenically cooled triple resonance probe. Resonance assignments were obtained from the following spectra: 2D ^1^H-^15^N HSQC, 2D ^1^H-^13^C HSQC, 3D HNCACB, 3D CBCA(CO)NH, 3D HNCO, 3D HBHA(CO)NH and 4D HCC(CO)NH-TOCSY^48^ spectra. All 3D and 4D spectra were acquired using non-uniform sampling and reconstructed using the maximum entropy algorithm in the Rowland NMR Toolkit^55^. ^13^C-aliphatic, ^13^C-aromatic and ^15^N-edited NOESY-HSQC spectra were acquired using uniform sampling for the extraction of interproton distance restraints.

Resonance assignments and integration of NOESY peaks were achieved using SPARKY^56^, followed by automatic peak list assignment, extraction of distance restraints, and structure calculation using the torsion angle dynamics package CYANA 3.0^57^. Dihedral-angle restraints derived from TALOS chemical shift analysis^58^ were also integrated in the calculation, with the restraint range set to twice the estimated standard deviation. CYANA was used to calculate 200 structures from random starting conformations then the best 20 structures were selected based on their stereochemical quality as judged using MolProbity^59^ (Supplementary Table S2).

### Homology modelling

Superfamilies that were identified to have a structural counterpart in the Protein Data Bank (see Supplementary Table S3) were modelled using the automodel parameters in Modeller v9.12^60^. A total of 100–500 models were generated for each structure. The lowest-energy models were selected and visualized using PyMOL v.2.0 (Pymol Molecular Graphics system, Schödinger, LLC).

### Phylogenetic analyses

We used Bayesian inference of phylogeny and maximum likelihood reconstruction to examine the molecular evolution of spider-venom ICK toxins. In order to search for non-venom outgroups, we examined datasets from one species of whip scorpion (*Mastigoproctus giganteus* (Uropygi); SRR1145698), which is non-venomous and regarded as the sister group to spiders^61^, and non-venom gland transcriptomes from three species of taxonomically diverse spiders: hypodermal tissue from *Cupiennius saliei* (SRR880446); leg tissue from *Liphistius malayanus* (SRR1145736); and leg tissue from *Neoscona arabesca* (SRR1145741). Transcriptomes were downloaded and converted to fastq files using fastq-dump v2.5x included in the SRA Toolkit (https://trace.ncbi.nlm.nih.gov/Traces/sra/sra.cgi?view=software), trimmed using trimmomatic^62^ with window quality 30, window size 4, minimum read length 60 bp, then assembled with Trinity^63^ using default settings. The resulting assemblies were combined and all predicted coding DNA sequences (CDSs) extracted using the “Get open reading frames (ORFs) or coding sequences (CDSs)” tool in Galaxy^64^. We then used BLAST+^65,66^ to search all members of each ICK superfamily against the resulting assemblies, as well as UniProtKB and the NCBI non-redundant (nr) and transcriptome shotgun assembled (TSA) databases, with an expected cut-off value of e-3.

The resulting hits were filtered based on the presence of a predicted signal peptide using SignalP^34^, and aligned with MAFFT by using L-INS-i^67^. The alignment was then adjusted manually in CLC Main Workbench v7.6.1 (CLC-Bio, Denmark) to correct misaligned structurally equivalent cysteine residues before realigning the inter-cysteine loops using the MAFFT regional re-alignment script (http://mafft.cbrc.jp/alignment/software/regionalrealignment.html). The resulting alignment was used for Bayesian inference of phylogeny by MrBayes v3.2.2^68^; we performed two simultaneous runs for 10,000,000 generations using lset rates=gamma with prset aamo-delpr = mixed command, which enables the program to optimize between nine different amino acid substitution matrices implemented in MrBayes. The log-likelihood score of each saved tree was plotted against the number of generations to establish the point at which the log likelihood scores reached their asymptote, and the posterior probabilities for clades was established by constructing a majority-rule consensus tree for all trees generated after completed burn-in phase. We also used IQ-Tree v1.3.6^69^ for phylogenetic reconstruction by maximum likelihood, using the TESTNEW command to let IQ-Tree determine the best fitting gene model according to a Bayesian information criterion analysis. Node support was tested by ultrafast bootstrap approximation (UFBoot)^70^ using 10,000 iterations. Trees were rooted using *Mastigoproctus giganteus* as an outgroup, and visualised and exported using Archaeopteryx v0.99 (http://www.phylosoft.org/archaeopteryx) and FigTree v1.3.1 (http://tree.bio.ed.ac.uk/software/figtree/).

## Supporting information

## Acknowledgments

We acknowledge financial support from the Australian Research Council (Discovery Grant DP130103813 to G.F.K. and DECRA Fellowship DE160101142 to E.A.B.U.) and the Australian National Health & Medical Research Council (Principal Research Fellowship to G.F.K.). We thank David Wilson for collecting *H. infensa* specimens, the Australian Genome Research Facility for bioinformatic assistance, Philip Lawrence for assistance with mass matching of MS data, and Julie Klint, Niraj Bende and Jessie Er for help with recombinant peptide production. We thank Michael Nuhn and Lien Li from the Bioinformatics Resource Australia-EMBL for help with data submission to the European Nucleotide Archive.

## Author contributions

S.S.P. reared and dissected spiders, generated and analysed cDNA libraries, designed, produced and purified recombinant peptides. S.S. produced and purified recombinant peptides. Y.K.Y.C. analysed NMR data and solved the reported DRP structures. M.M. helped with NMR data acquisition. E.A.B.U. performed the phylogenetic analysis. C.D., P.E., E.A.B.U. and G.M.N., acquired and analysed MS data. Q.K. assisted with bioinformatic analysis. S.S.P., Y.K.Y.C., E.A.B.U. and G.F.K. wrote the manuscript. G.F.K. conceived the research, with input from J.S.M., and provided funding and facilities. All authors revised the manuscript.

## Data deposition

Metadata and annotated nucleotide sequences generated for the *Hadronyche infensa* venom-gland transcriptome were deposited in the European Nucleotide Archive under project accession number ERA298588 (HACE01000001-HACE0100120). Atomic coordinates of peptide structures solved in this study were deposited in the Protein Data Bank under accession codes 2N6N, 2N6R, 6BA3 and 2N8K, while corresponding NMR chemical shifts were deposited in BioMagResBank under accession numbers BMRB27774, BMRB25778, BMRB25853 and BMRB30352.

