## Supplementary material for "Structural venomics: evolution of a complex chemical arsenal by massive duplication and neofunctionalization of a single ancestral fold"

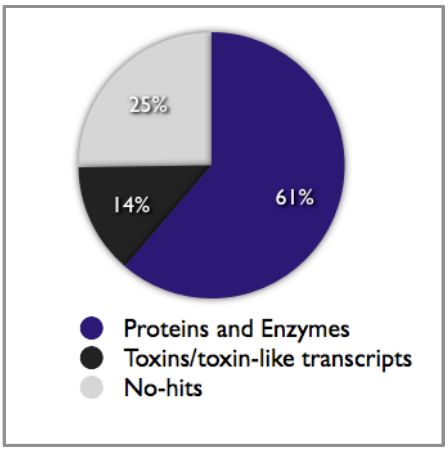


**Figure S1:** Categories of ESTs obtained from 454 sequencing of a cDNA library generated using venom-glands from several *Hadronyche infensa* specimens.

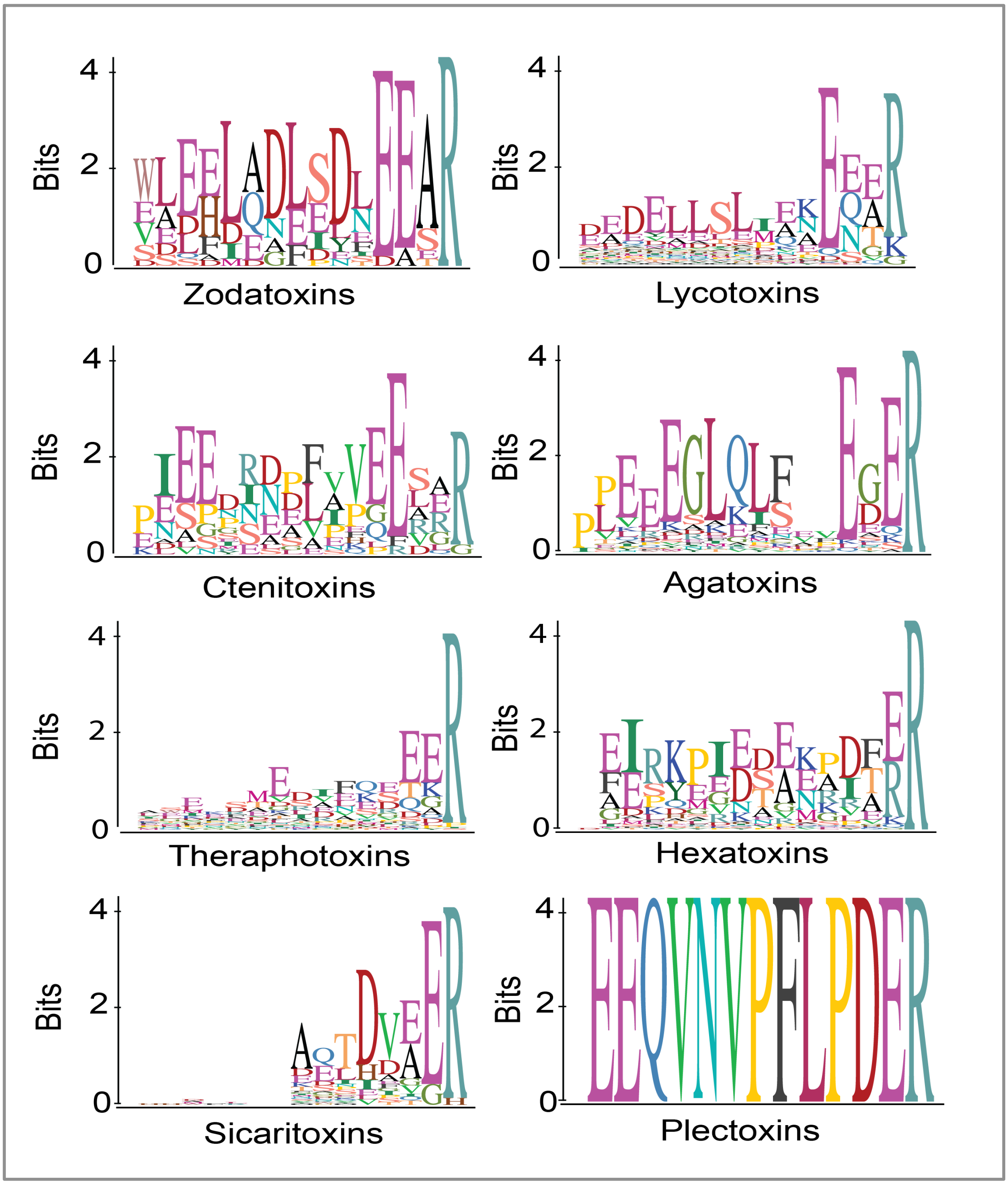


**Figure S2:** Sequence Logo analysis of the propeptide regions of precursors encoding the following families of spider-venom peptides: hexatoxins, lycotoxins, agatoxins, zodatoxins, ctenitoxins and theraphotoxins. (Data were obtained in 2008 obtained by David Wood from all available transcript information present on ArachnoServer (www.arachnoserver.org) before the beginning of this study.)

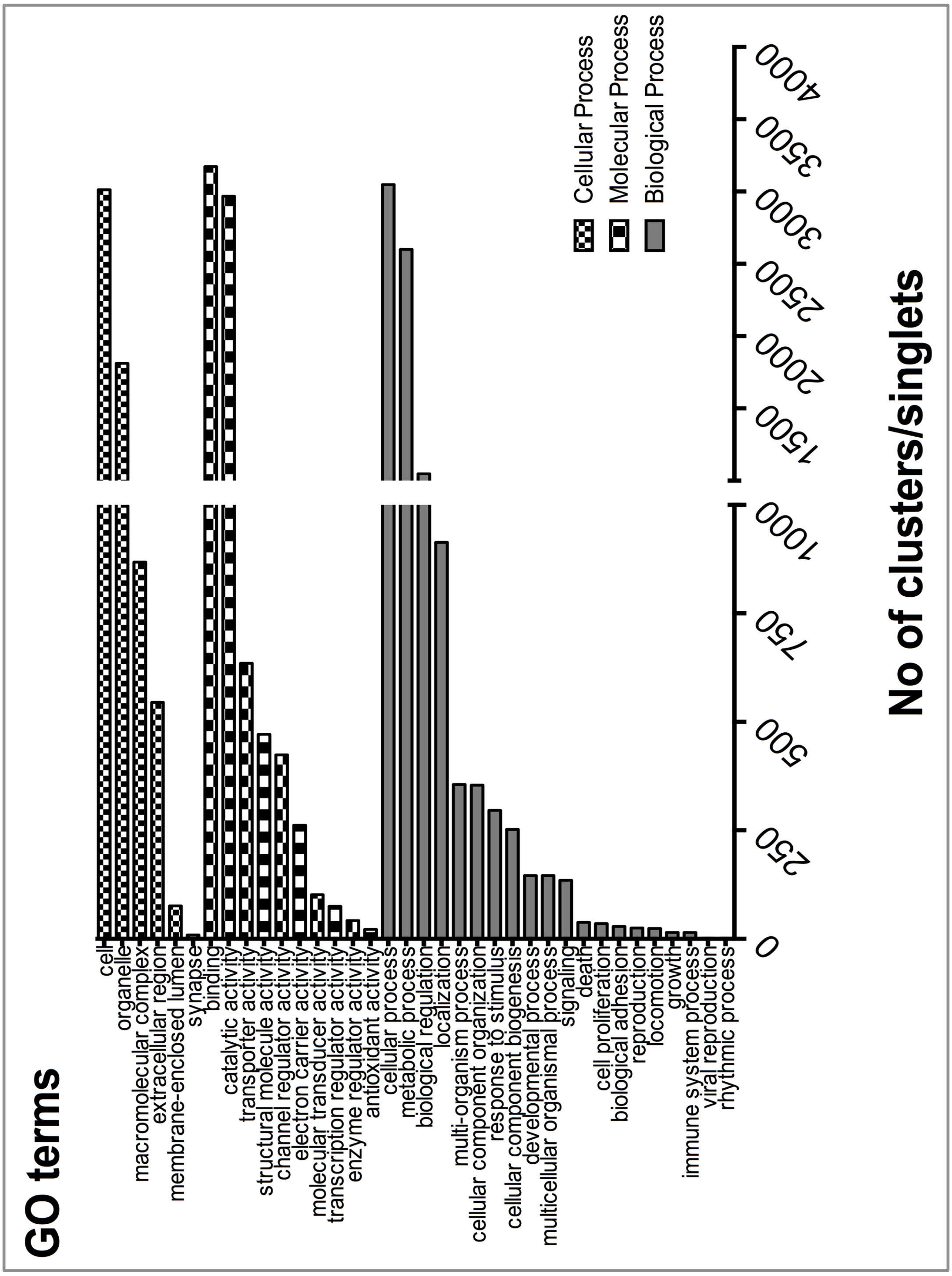


**Figure S3:** GO distribution by level of venom-gland ESTs from *H. infensa*. This representation shows the number of sequences at the nodes of level 2 of the Gene Ontology (GO) distribution for the three different tree types: Biological, molecular and cellular processes.

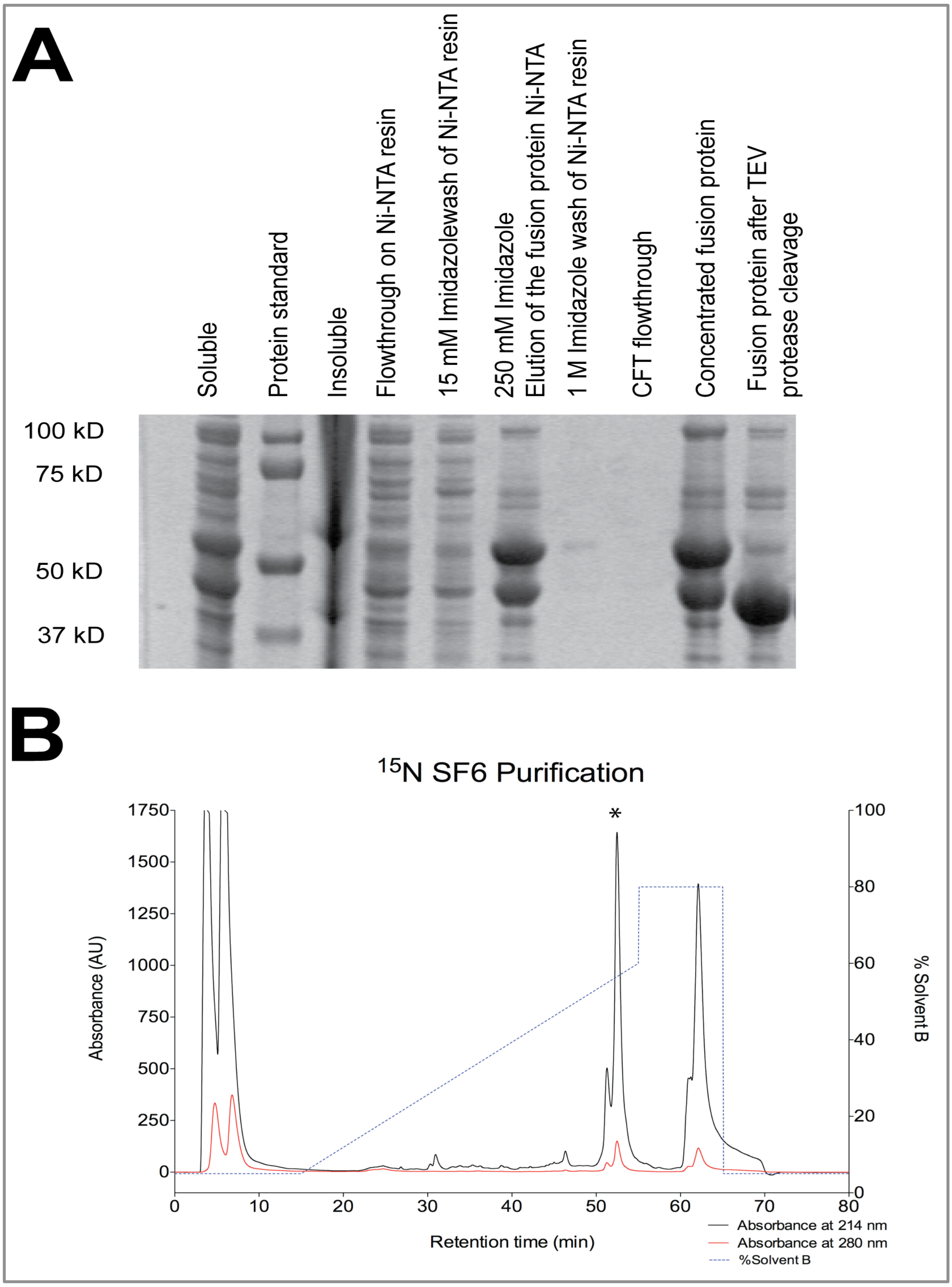


**Figure S4:** (A) SDS-PAGE gel and (B) RP-HPLC chromatogram summarizing the expression and purification of $^{15}$N-labelled SF6 peptide. Gel lanes are as follows: “Soluble” and “insoluble” fractions following cell lysis; flow-through from addition of soluble cell fraction to Ni-NTA resin; eluates obtained from washing Ni-NTA resin with 15 mM, 250 mM and 1 M imidazole; MBP-toxin fusion protein before and after cleavage with tobacco etch virus (TEV) protease. The peak marked with an asterisk in the RP-HPLC chromatogram corresponds to the peptide with the expected mass. The ordinate axis indicates the gradient in percentage acetonitrile (solvent B).

**Table S1:** Transcript abundance (TPMs) for each toxin superfamily identified in the *Hadronyche infensa* venom-gland transcriptome.

| Superfamily | Superfamily Name | TPMs |
|---|---|---|
| 1 | Ah Puch | 342.02 |
| 2 | Anat | 11917.8 |
| 3 | Hades | 12893.8 |
| 4 | Erlik | 9660.52 |
| 5 | Chi You | 0 |
| 6 | ‘Oro | 456.35 |
| 7 | Pele | 2343.58 |
| 8 | Ixtab | 636.87 |
| 9 | Shiva | 2875.08 |
| 10 | Kisin | 5624.89 |
| 11 | Thanatos | 2.43 |
| 12 | Sekhmet | 1088.15 |
| 13 | Ares | 9556.72 |
| 14 | Wurrukatte | 312.99 |
| 15 | Pluto | 16.19 |
| 16 | Zahhak | 140.73 |
| 17 | Nergal | 1226.1 |
| 18 | Grim reaper | 14.14 |
| 19 | Yama | 1898.66 |
| 20 | Ereshkigal | 2024.97 |
| 21 | Anubis | 6.82 |
| 22 | Loke | 55.77 |
| 23 | Qamaits | 9064.72 |
| 24 | Ankou | 270.63 |
| 25 | Tezcatlipoca | 11.52 |
| 26 | Set | 5101.63 |
| 27 | Mara | 33.51 |
| 28 | Tuoni | 31.05 |
| 29 | Mars | 87.6 |
| 30 | Sauron | 60.84 |
| 31 | Hulda | 47.44 |
| 32 | Midgardsormen | 15.43 |
| 33 | Menhit | 14 |

**Table S2:** Structural statistics for the NMR ensembles of SF6, SF22, SF23 and SF26 toxins.[a]

| Toxin family | **SF6** | **SF22** | **SF23** | **SF26** |
|---|---|---|---|---|
| PDB ID | 2N6R | 2N8K | 2N6N | 6BA3 |
| Experimental restraints | | | | |
| Inter-proton distance restraints | | | | |
| Total | 1657 | 1369 | 809 | 1099 |
| Intra-residue (*i* = *j*) | 375 | 336 | 118 | 293 |
| Sequential (\|*i* - *j*\| = 1) | 470 | 464 | 107 | 333 |
| Medium range (1 < \|*i* – *j*\| < 5) | 309 | 170 | 74 | 169 |
| Long range (\|*i* – *j*\| ≥ 5) | 503 | 399 | 116 | 304 |
| Disulfide bond restraints | 12 | 15 | 12 | 12 |
| Dihedral-angle restraints (ϕ, ψ) | 113 | 112 | 56 | 149 |
| ϕ dihedral angle restraints | 55 | 54 | 28 | 61 |
| ψ dihedral angle restraints | 58 | 58 | 28 | 63 |
| Total number of restraints per residue | 23.4 | 19.7 | 27.4 | 25 |
| Violations of experimental restraints | 0 | 1 | 0 | 1 |
| RMSD from mean coordinate structure (Å)[b] | | | | |
| All backbone atoms | 0.41 ± 0.08 | 0.65 ± 0.17 | 0.46 ± 0.15 | 0.37 ± 0.07 |
| All heavy atoms | 1.03 ± 0.14 | 1.23 ± 0.17 | 1.19 ± 0.21 | 0.74 ± 0.05 |
| Backbone atoms (Non-flexible regions[c]) | 0.35 ± 0.07 | 0.47 ± 0.13 | 0.22 ± 0.12 | 0.26 ± 0.06 |
| Heavy atoms (Non-flexible regions) | 1.01 ± 0.14 | 1.04 ± 0.14 | 1.00 ± 0.19 | 0.67 ± 0.06 |
| Stereochemical quality[d] | | | | |
| Ramachandran plot statistics | | | | |
| Residues in most favored Ramachandran region (%) | 92.1 ± 1.4 | 86.4 ± 1.1 | 93.5 ± 2.0 | 95.0 ± 1.2 |
| Disallowed regions [%] | 0.0 ± 0.0 | 0.0 ± 0.0 | 0.0 ± 0.0 | 0.1 ± 0.4 |
| Unfavorable sidechain rotamers [%] | 11.5 ± 2.1 | 7.6 ± 1.7 | 0.7 ± 0.7 | 6.9 ± 2.3 |
| Clashscore, all atoms[e] | 0.0 ± 0.0 | 0.0 ± 0.0 | 0.0 ± 0.0 | 0.3 ± 0.4 |
| Overall MolProbity score | 1.78 ± 0.08 | 2.20 ± 0.07 | 1.17 ± 0.26 | 1.54 ± 0.17 |

[a] All statistics are given as mean ± S.D.

[b] Mean RMSD calculated over the entire ensemble of 20 structures.

[c] The non-flexible region for each peptide is defined as follows:

SF6: Residues 2–74

SF22: Residues 4–17, 26–76

SF23: Residues 4–28

SF26: Residues 2–71

[d] Sterochemical quality according to MolProbity (http://helix.research.duhs.duke.edu).

[e] Clashscore is defined the number of steric overlaps >0.4 Å per 1000 atoms.

**Table S3:** List of homology models generated for this study.

| Superfamily | Protein Data Bank ID used for homology modelling |
|---|---|
| SF2 | 2KRA |
| SF5 | 1KFP |
| SF8 | 1KOZ |
| SF9 | 1AXH |
| SF10 | 1G9P |
| SF11 | 3L0R |
| SF13 | 1VTX |
| SF17 | 1S6X |
| SF20 | 2H1Z |
| SF22 | 2NK8 |
| SF23 | 2N6N |
| SF27 | 1FCU |
| SF28 | 1POC |
| SF29 | 1QNX |
| SF32 | 1JC9 |

#### Summary of accession numbers used in this study

##### Sequence read archive accessions:

*Cupiennius saliei* (SRR880446); *Liphistius malayanus* (SRR1145736); *Neoscona arabesca* (SRR1145741) and *Mastigoproctus giganteus* (SRR1145698).

The species abbreviations shown in parentheses below are as follows: *Apomastus schelingeri*: APOSC; *Brachypelma smithi*: BRASM; *Chilobrachys guangxiensis*: CHIGU; *Hadronyche infensa*: HADIN/Hi; *Hadronyche modesta*: HADMO; *Hadronyche robustus*: ATRRO; *Hadronyche venenata*: HADVN; *Hadronyche versuta*: HADVE; *Haplopelma schmidti*: HAPHA, HAPSC; *Lassiodora sp.*: LASSB; *Macrothele gigas*: MACGS; *Trittame loki*: TRILK.

ABY77683, ACD01232, ACD01229, ABY77684, ABY77685, ABY77686, ACD01231, ACD01230, ACD01228, D2Y251 (H18A1_HAPHA), D2Y2P1 (H18G1_HAPA), D2Y2P2 (H18A2_HAPHA), D2Y2N9 (H18E1_HAPHA), D2Y2H1 (H18C1_HAPHA), D2Y252 (H18B1_HAPHA), D2Y2H2 (H18D1_HAPHA), D2Y2P0 (H18F1_HAPHA), B3FIN4 (TX18A_HAPSC), B1P1I3 (JTZ64_CHIGU), B1P1I4 (JTZ65_CHIGU), BAM15662, BAN13533, ADF28499, W4VS46 (ICK10_TRILK), W4VRV2 (ICK11_TRILK), W4VSI8 (ICK8_TRILK), W4VSB9 (ICK9_TRILK), W4VRX0 (ICK22_TRILK), W4VSB0 (ICK24_TRILK), W4VSI5 (ICK23_TRILK), Q75WG7 (TXM12_MACGS), Q9BJV8 (TOT2A_ATRIL), Q9BJV7 (TOT2B_ATRIL), Q9BJV9 (TOT2A_HADIN), Q9BJW0 (TOT2B_HADIN), BAN13534, B3FIP1 (TZ721_HAPSC), B3FIP2 (TZ722_HAPSC), B1P1J5 (TZ72_CHIGU), W4VRX8 (ICK12_TRILK), W4VRU3 (ICK41_TRILK), W4VRV7 (ICK42_TRILK), W4VSI7 (ICK13_TRILK), W4VS32 (ICK15_TRILK), W4VSB2 (ICK19_TRILK), W4VSB7 (ICK14_TRILK), W4VRV1 (ICK16_TRILK), W4VRX3 (ICK17_TRILK), P49267 (TXP1_APOSC), P49269 (TXP4_APOSC), P49270 (TXP6_APOSC), P499272 (TXP9_APOSC), W4VSI6 (ICK18_TRILK), B3FIV1 (TXBS1_BRASM), A3F7X1 (TXLT4_LASSB), ADF28491, Q75WH1 (TXPT6_MACGS), P83558 (TXMG_MACGS), W4VRV3 (ICK6_TRILK), W4VS20 (ICK25_TRILK), W4VRW4 (ICK32_TRILK), W4VRU8 (ICK26_TRILK), W4VRU5 (ICK36_TRILK), W4VRW1 (ICK37_TRILK), W4VRY7 (ICK7_TRILK), W4VRY9 (ICK2_TRILK), W4VRW6 (ICK27_TRILK), W4VRU6 (ICK31_TRILK), W4VS15 (ICK30_TRILK), W4VSI4 (ICK28_TRILK), W4VSA7 (ICK29_TRILK), W4VSI3 (ICK33_TRILK), W4VSA5 (ICK34_TRILK), W4VS12 (ICK35_TRILK), W4VSA2 (ICK39_TRILK), W4VSI1 (ICK38_TRILK), W4VS08 (ICK40_TRILK), B1P1H6 (JZT56_CHIGU), B1P1H7 (JZT57_CHIGU), CDF44148, S0F1N7 (TO1B_HADVN), CDF44149, CDF44150, S0F1M4 (TO1A_HADVN), CDF44153, A5A3H4 (TO1E_ATRRO), A5A3H3 (TO1D_ATRRO), A5A3H5 (TO1F_ATRRO), A5A3H2 (TO1C_ATRRO), P0DMQ1 (TO1B_HADMO), P0DMQ0 (TO1A_HADMO), P0DMQ2 (TO1C_HADMO), S0F1N6 (TO1D_HADIN), S0F1N0 (TO1E_HADIN), S0F204 (TO1F_HADIN), CDF44165, S0F215 (TO1G_HADIN), CDF44166, P83580 (TO1A_ATRRO), A5A3H1 (TO1B_ATRRO*)*, CDF44155, CDF44163, CDF44164, CDF44170, S0F1M6 (TOK1G_ATRRO), S0F209 (TOK1H_HADVE), P83560 (TXMG4_MACGS), Q75WG5 (TXM14_MACGS)

*Haplopelma schmidti* clade:

ABY77692, ABY77701, ABY77698, ABY77695, ABY77694, ABY77697, ABY77699, ABY77696, D2Y274 (H16E1_HAPHA), ABY77693, D2Y262 (H1610_HAPHA), D2Y263 (H1611_HAPHA), D2Y253 (H16A1_HAPHA), D2Y271 (H16B1_HAPHA), D2Y272 (H16C1_HAPHA), D2Y277 (H16G1_HAPHA), D2Y297 (H1624_HAPHA), D2Y276 (H16F2_HAPHA), D2Y287 (H16Q1_HAPHA), D2Y254 (H16A2_HAPHA), D2Y281 (H16K1_HAPHA), D2Y267 (H1615_HAPHA), D2Y275 (H16F1_HAPHA), D2Y298 (H1625_HAPHA), D2Y2I2 (H1627_HAPHA), D2Y2I0 (H16X1_HAPHA), D2Y2I1 (H1626_HAPHA), D2Y264 (H1612_HAPHA), D2Y260 (H16A8_HAPHA), D2Y261 (H16A9_HAPHA), D2Y286 (H16P1_HAPHA), D2Y295 (H1622_HAPHA), D2Y2P8 (H16B2_HAPHA), D2Y2P6 (H1629_HAPHA), D2Y2P3 (H1628_HAPHA), D2Y2P5 (H16ZA_HAPHA), D2Y293 (H1620_HAPHA), D2Y280 (H16J1_HAPHA), D2Y268 (H1616_HAPHA), D2Y294 (H1621_HAPHA), D2Y288 (H16R1_HAPHA), D2Y253 (H16A6_HAPHA), D2Y259 (H16A7_HAPHA), D2Y265 (H1613_HAPHA), D2Y255 (H16A3_HAPHA), D2Y2H9 (H16V1_HAPHA), D2Y270 (H1618_HAPHA), D2Y292 (H1619_HAPHA), D2Y257 (H16A5_HAPHA), D2Y291 (H16U1_HAPHA), D2Y269 (H1617_HAPHA), D2Y290 (H16T1_HAPHA), D2Y289 (H16S1_HAPHA), D2Y284 (H16N1_HAPHA), D2Y285 (H16O1_HAPHA), D2Y296 (H1623_HAPHA), D2Y256 (H16A4_HAPHA), D2Y266 (H1614_HAPHA), D2Y279 (H16I1_HAPHA), D2Y278 (H16H1_HAPHA), D2Y273 (H16D1_HAPHA), ABY77702, ABY77703, D2Y283 (H16M1_HAPHA), D2Y299 (H19A1_HAPHA,) B3FIQ7 (TX16C_HAPSC).

### Superfamily diversity

#### Peptides smaller than 16 kDa

##### Superfamily 1 [Ah Puch]

Transcripts belonging to Superfamily 1 (SF1) encode 128-residue prepropeptides. The mature peptide spans 75 amino acid residues and includes 12 cysteines that form six disulfide bonds (Figure 1.1a). Remarkably, the mature toxin contains an internal repeat of two ICK-like sequences, a rare but documented phenomenon has been noted in other spider species.[1,2] For example, the central region of the first repeat, which comprises residues 48–88 of the precursor, contains the sequence "**CCE**G**LECW**K**RR**" whereas the second repeat has the sequence "**CCE**E**LECW**E**RR**" (residues that are identical in both repeats are shown in bold). The two ICK domains show a high level of sequence identity with PcTx1 (62% and 50% for the N- and C-terminal ICK domains, respectively) from the Trinidad chevron tarantula *Psalmopoeus cambridgei*. PcTx1 is the most potent and selective known inhibitor of acid-sensing ion channel 1a (ASIC1a); it blocks the rat channel with an $IC_{50}$ of ~1 nM. Previous work in our lab identified the pharmacophore of PcTx1[3] and many of these functional residues are conserved in Superfamily 1 peptides.

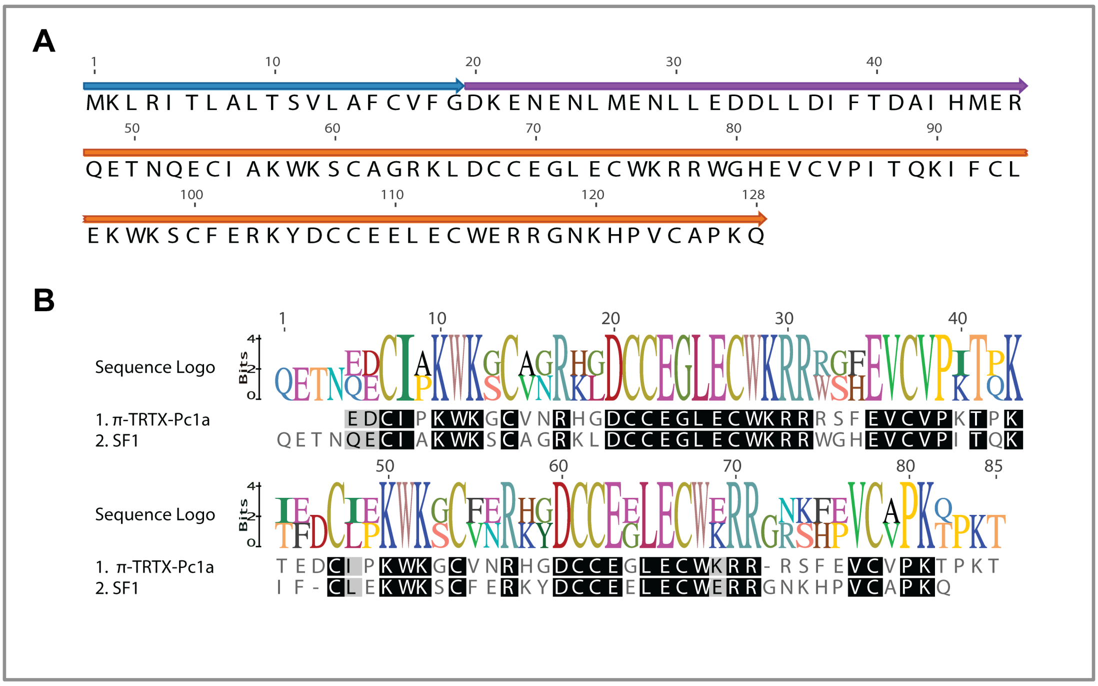


**Figure 1.1:** (a) Schematic of the 128-residue precursor encoding the SF1 toxin π-hexatoxin-Hi1a. The signal peptide, propeptide, and mature toxin are shown in blue, purple, and orange, respectively. (b) Sequence alignment showing amino acid identities (boxed in black) between PcTx1 and the two repeated domains within π-hexatoxin-Hi1a.

Thus, we surmised that SF1 peptides might block ASIC1a. The sequence shown in Figure 1.1a was made recombinantly and shown to block rat ASIC1a with an $IC_{50}$ of 0.40 nM and human ASIC1a with an 0.52 nM[4]. This peptide is therefore the most potent blocker of ASIC1a discovered to date. In line with the rational nomenclature developed for naming spider-venom peptides[5], we named this peptide π-hexatoxin-

Hi1a, where the π prefix indicates activity on ASICs.

#### Superfamily 2 [Anat]

Superfamily 2 peptides are expressed as 90-residue precursors that lack a propeptide region (Figure 1.2a). The mature peptide spans 68 residues with a C-terminal amidation signal. These peptides correspond to the MIT-like atracotoxins that were initially isolated from the venom of the Blue Mountains funnel-web spider *Hadronyche versuta* [6]. These peptides are now known to be part of the AVIT superfamily of peptides that includes mamba intestinal toxin 1 (MIT1) from the venom of the black mamba snake, Bv8 and orthologs from skin secretions of toads of the genus *Bombina*, and the human cytokine-like prokineticins[7].

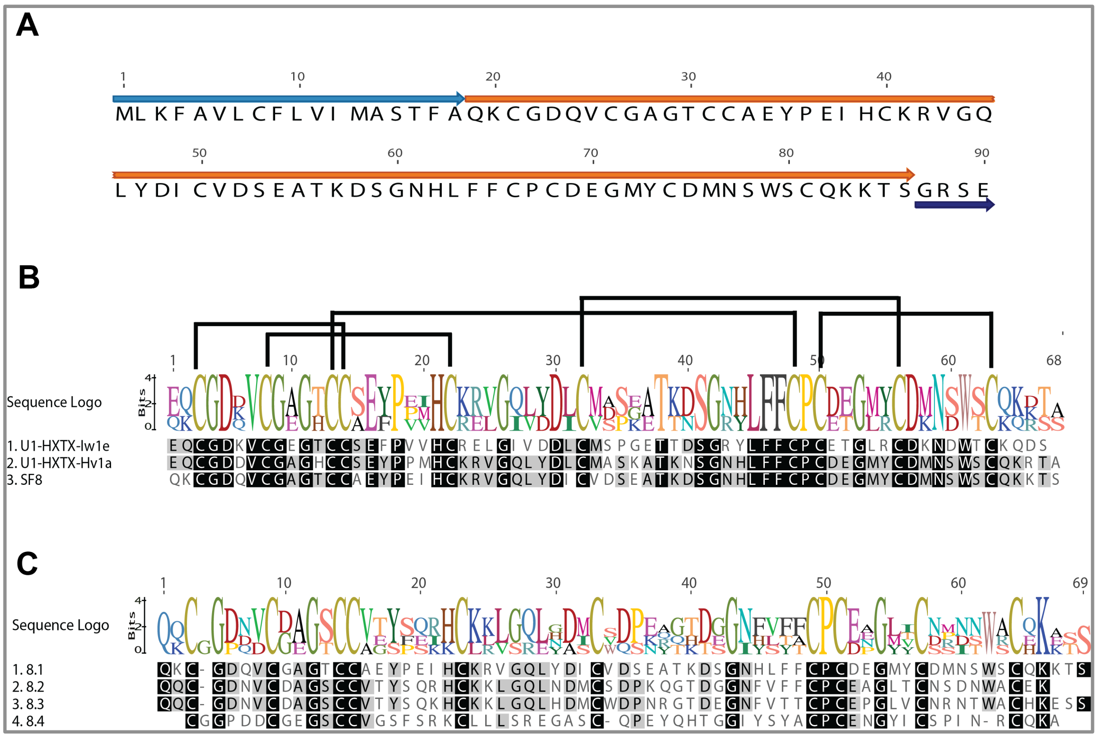


**Figure 1.2:** (a) Schematic of 99-residue precursor encoding Superfamily 2 peptide U$_1$-hexatoxin-Hi1a. The signal peptide, mature toxin, and C-terminal amidation signal are shown in light blue, orange and dark blue, respectively. (b) Sequence alignment showing amino acid identities (boxed in black) between U$_1$-hexatoxin-Hi1a and MIT-like peptides from *Illawara wisharti* (Iw) and *H. versuta* (Hv). The disulfide-bond framework is shown above the sequence alignment. (c) Alignment of Superfamily 2 (SF2) isoforms from *H. infensa*.

Unlike other members of the AVIT superfamily, the spider toxins lack the N-terminal "AVIT" sequence and do not display any of the activities reported for other family members (Figure 1.2a). Thus, while the 3D structure of these spider toxins is likely very similar to that of MIT1[8] and Bv8[9], their function in venom remains enigmatic. Figure 1.2b also shows the sequence alignment of other MIT-like peptides isolated from the Australian funnel-web spiders *H. versuta* and *Illawara wisharti,* while Figure 1.2c shows the alignment of some Superfamily 2 (SF2) isoforms (paralogs) from *H. infensa*.

#### Superfamily 3 [Hades]

Superfamily 3 peptides are expressed as 83-residue prepropeptides, with the mature toxin spanning 38 residues (Figure 1.3a). The prototypic family member $U_2$-hexatoxin-Hi1a (formerly known ACTX-HiOB4219) includes eight cysteines arranged in four disulfide bonds, three of which form an ICK motif.

The structure of $U_2$-hexatoxin-Hi1a solved using homonuclear NMR spectroscopy revealed two different conformations of equal abundance due to *cis-trans* isomerisation of Pro30[10]. It was suggested (but not shown experimentally) that this family of peptides are insecticidal based on limited sequence and structural similarities with the insecticidal μ-agatoxin peptides isolated from the venom of the unrelated American funnel-web spider *Agelena orientalis*[10] (Figure 1.3b).

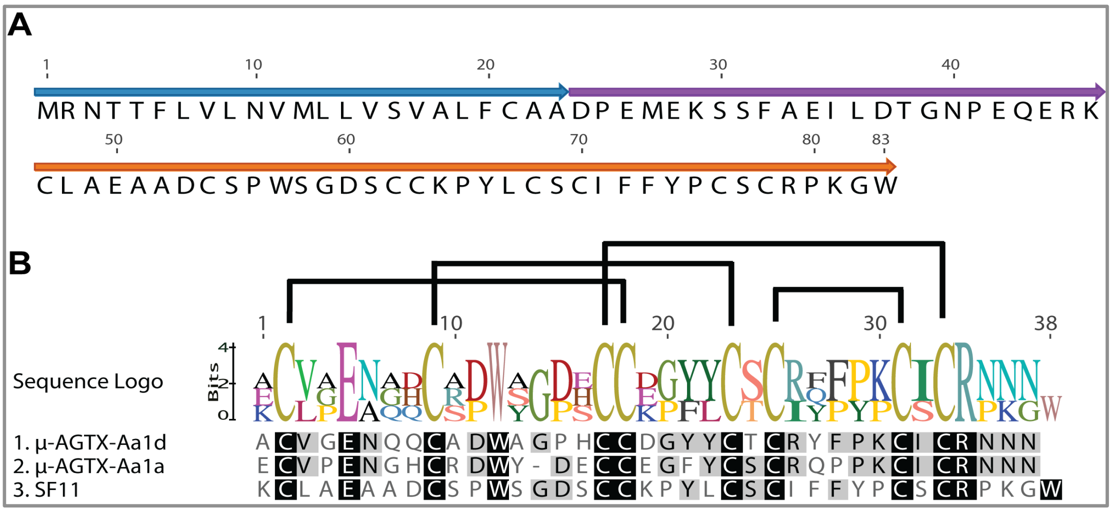


**Figure 1.3:** (a) Schematic of 83-residue precursor encoding the Superfamily 3 peptide $U_2$-hexatoxin-Hi1a. The signal peptide, propeptide, and mature toxin are shown in blue, purple, and orange, respectively. (b) Sequence alignment showing amino acid identities (boxed in black) between μ-Aga-I and μ-Aga-IV from the venom of the American funnel-web spider *A. orientalis* and the Superfamily 3 peptide $U_2$-hexatoxin-Hi1a; the disulfide-bond pattern is shown above the sequence logo.

#### Superfamily 4 [Erlik]

Superfamily 4 peptides are expressed as ~120-residue prepropeptide precursors with the mature toxin spanning 69 residues (Figure 1.4a). The prototypic family member $U_3$-hexatoxin-Hi1a is an ortholog of $U_7$-theraphotoxin-Hs3a from the Chinese bird spider *Haplopelma schmidti* and $U_{15}$-hexatoxin-Mg1b from the hexathelid spider *Macrothele gigas* (Figure 1.4b). The transcript encoding this toxin was the most abundant in the venom-gland transcriptome; this high abundance combined with the fact that the toxin is taxonomically well conserved, at least within mygalomorph spiders, and that insecticidal activity has been demonstrated for at least one superfamily member ($U_{15}$-hexatoxin-Mg1b)[11], suggests that these toxins are an important component of the insecticidal toxin arsenal of these spiders. The disulfide scaffold in these toxins is unique with a large gap between cysteines 7 and 8. Hence, SF4 toxins may have a novel 3D fold.

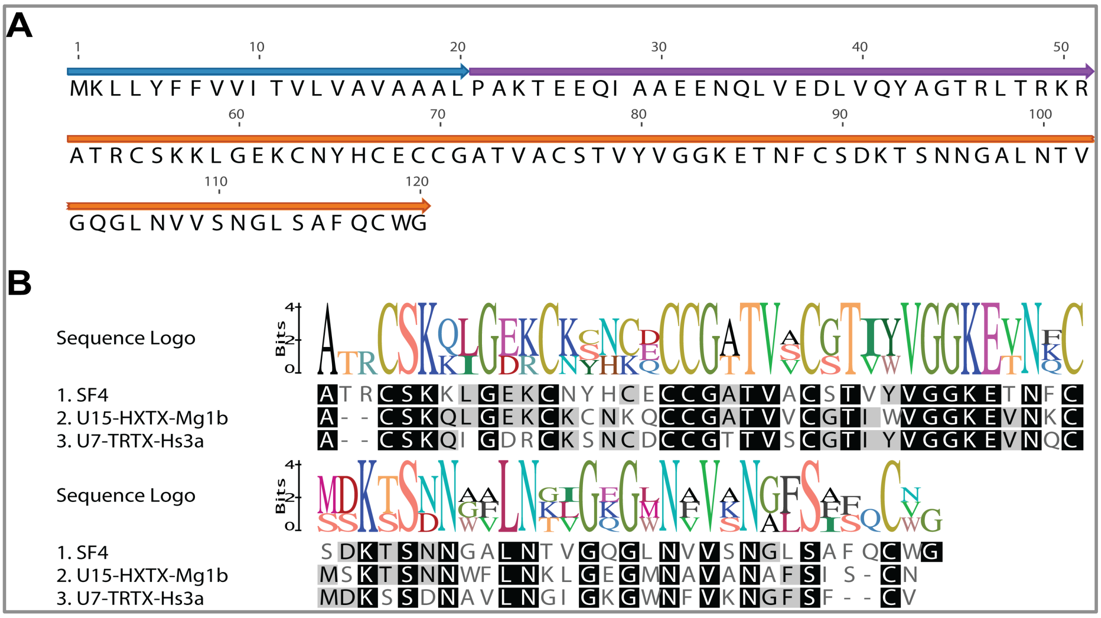


**Figure 1.4:** (a) Schematic of the 120-residue precursor encoding the Superfamily 4 peptide $U_3$-hexatoxin-Hi1a. The signal peptide, propeptide, and mature toxin are shown in blue, purple, and orange, respectively. (b) Sequence alignment showing amino acid identities (boxed in black) between $U_7$-theraphotoxin-Hs3a, $U_{15}$-hexatoxin-Mg1b and $U_3$-hexatoxin-Hi1a.

#### Superfamily 5 [Chi You]

Superfamily 5 peptides are expressed as ~84-residue prepropeptides, with the mature peptide spanning 18 amino acid residues (Figure 1.5a). These sequences are clearly orthologs of gomesin (Figure 1.5b) an antimicrobial peptide isolated from hemocytes of the spider *Acanthoscurria gomesiana*. Hemocytes are an important part of the immune system of invertebrates, and they can be likened to vertebrate phagocytes. Gomesin has an amidated C-terminus and an N-terminal pyroglutamate (pyroGlu) residue. The *H. infensa* peptide nominally begins with an N-terminal glutamine residue, as does gomesin, and we presume that like gomesin it spontaneously cyclises to yield pyroGlu.

The precursor of the *H. infensa* gomesin-like peptide also contains a "KR" amidation signal immediately downstream of the final Arg residue and thus we predict that it is C-terminally amidated like gomesin. Gomesin forms a two-stranded antiparallel β sheet, connected by a non-canonical β turn, and it includes four cysteines that form two disulfide bonds[12]. Gomesin has broad-range antimicrobial activity and it is assumed to be part of the inmate immune response of spiders. This raises the question of whether the gomesin-like peptide expressed in the venom of *H. infensa* is antimicrobial and, if so, what is its role in the venom?

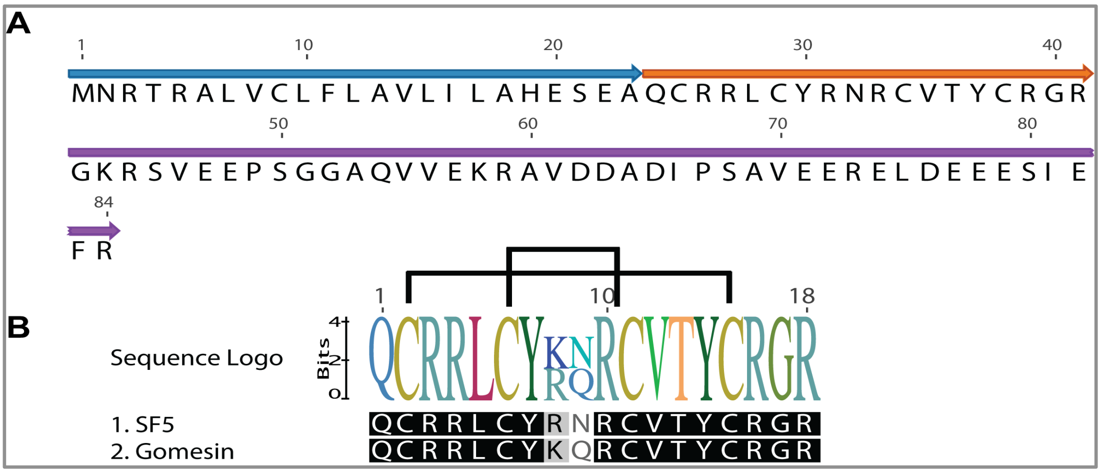


**Figure 1.5:** (a) Schematic of the 84-residue precursor encoding the Superfamily 5 gomesin-like peptides. The signal peptide, propeptide, and mature toxin are shown in blue, purple, and orange, respectively. (b) Sequence alignment showing amino acid identities (boxed in black) between the gomesin-like peptide expressed in the venom gland of *H. infensa* and gomesin from hemocytes of *Acanthoscurria gomesiana*. Disulfide bonds are indicated above the sequence logo.

#### Superfamily 6 [‘ORO]

Superfamily 6 peptides are expressed as 94-residue precursors that lack a propeptide region; the precursor of the prototypic family member $U_4$-hexatoxin-Hi1a encodes an 18-residue signal peptide and a 76-residue mature toxin (Figure 1.6a). These toxins are orthologs of toxins previously isolated from the Chinese tarantulas *Haplopelma schmidti* and *Chilobrachys guanxiensis*, both with unknown target and 3D structure (Figure 1.6b).

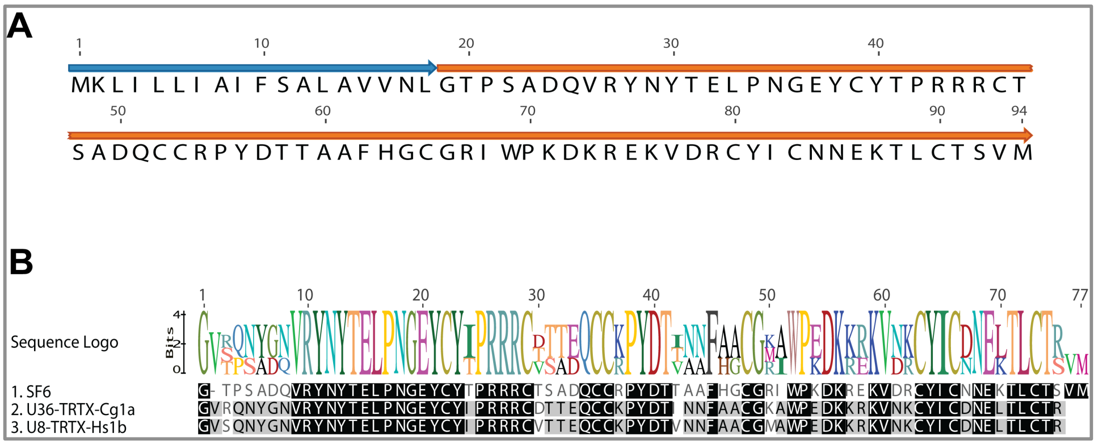


**Figure 1.6:** (a) Schematic of 94-residue precursor encoding the Superfamily 6 peptide $U_4$-hexatoxin-Hi1a. The signal peptide and mature toxin are shown in blue and orange, respectively. (b) Sequence alignment showing amino acid identities (boxed in black) between $U_8$-theraphotoxin-Hs1b, $U_{36}$-theraphotoxin-Cg1a, and $U_4$-hexatoxin-Hi1a.

#### Superfamily 7 [Pele]

Superfamily 7 peptides are expressed as 148-residue prepropeptide precursors, with the mature peptide spanning 85 residues (Figure 1.7a). A BLAST search revealed significant homology between the prototypic family member $U_5$-hexatoxin-Hi1a and $U_3$-theraphotoxin-Pm1a from the king baboon spider *Pelinobius muticus* (formerly known as *Citharischius crawshayi*) (Figure 1.7b). This peptide is unusual as it contains an odd number of cysteines and its cleavage sites are difficult to predict. However, we predicted the most likely propeptide cleavage site to be after position 38; it is also unclear whether the large Pro/Thr-rich C-terminal "tail" after the last cysteine residue is post-translationally modified.

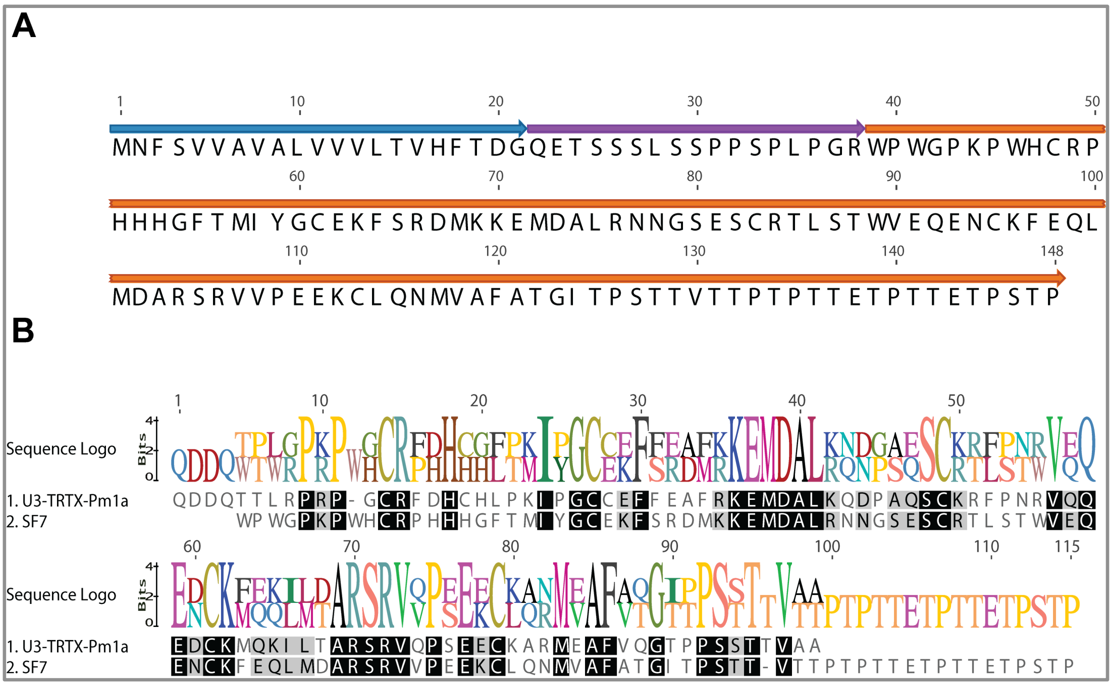


**Figure 1.7:** (a) Schematic of 148-residue precursor encoding the Superfamily 7 peptide $U_5$-hexatoxin-Hi1a. The signal peptide, propeptide, and mature toxin are shown in blue, purple, and orange, respectively. (b) Sequence alignment showing amino acid identities (boxed in black) between $U_5$-hexatoxin-Hi1a and $U_3$-theraphotoxin-Pm1a from the King baboon spider.

#### Superfamily 8 [Ixtab]

Superfamily 8 peptides are expressed as 109-residue prepropeptide precursors. The mature toxin spans 42 residues and includes six cysteines that are predicted to form three disulfide bonds (Figure 1.8). The mature toxin has ~34% sequence identity with β-TRTX-Cj1a, an ICK toxin from the Chinese tarantula *Chilobrachys guanxiensis.* β-TRTX-Cj1a inhibits $Na_V1.5$ channels with an $IC_{50}$ of 380 nM and the $K_V2.1$ channel with an $IC_{50}$ of 430 nM[13,14]. Although Superfamily 8 peptides are likely to adopt a classical ICK fold as for β-TRTX-Cj1a, the relatively low level of sequence identity makes it difficult to predict their molecular target.

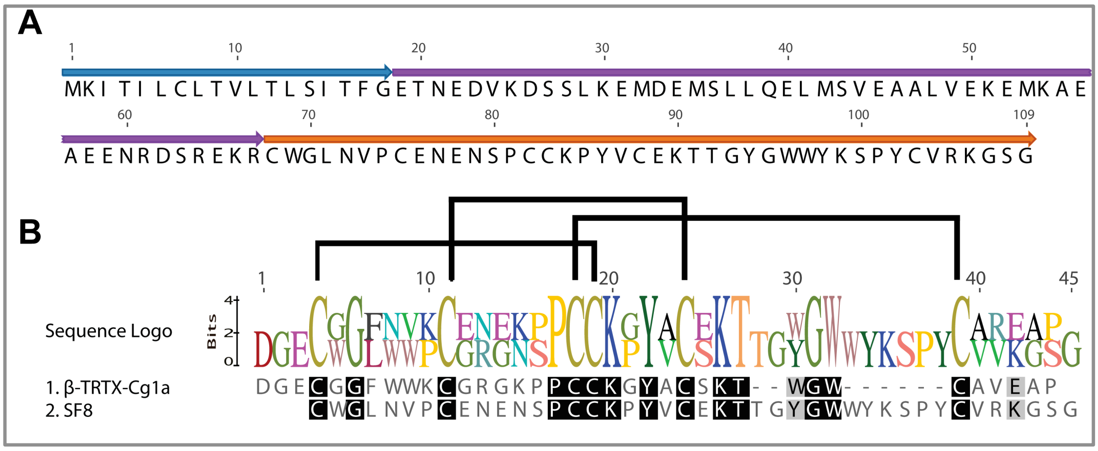


**Figure 1.8:** (a) Schematic of 109-residue precursor encoding the Superfamily 8 peptide $U_6$-hexatoxin-Hi1a. The signal peptide, propeptide, and mature toxin are shown in blue, purple, and orange, respectively. (b) Sequence alignment showing amino acid identities (boxed in black) between the mature toxin $U_6$-hexatoxin-Hi1a and β-TRTX-Cg1a from the Chinese tarantula *Chilobrachys guanxiensis*. Disulfide-bond connectivities are shown above the sequence logo.

#### Superfamily 9 [Shiva]

Superfamily 9 corresponds to the well-studied ω-hexatoxin-1 family[15-17]. These peptides are expressed as 78-residue prepropeptide precursors, with the mature toxin spanning 36–37 residues (Figure 1.9). These toxins were the first peptides isolated from venom of Australian funnel-web spiders and shown to be insecticidal. The mature toxin contains six conserved cysteines that are paired to form three disulfide bridges, which form an ICK motif[15]. The prototypic family member ω-hexatoxin-Hv1a blocks insect (but not vertebrate) $Ca_V$ channels[17,18]. However, it was subsequently shown that some members of this superfamily also block insect calcium-activated potassium ($K_{Ca}$) channels, as exemplified by κ-hexatoxin-Hv1c[19,20].

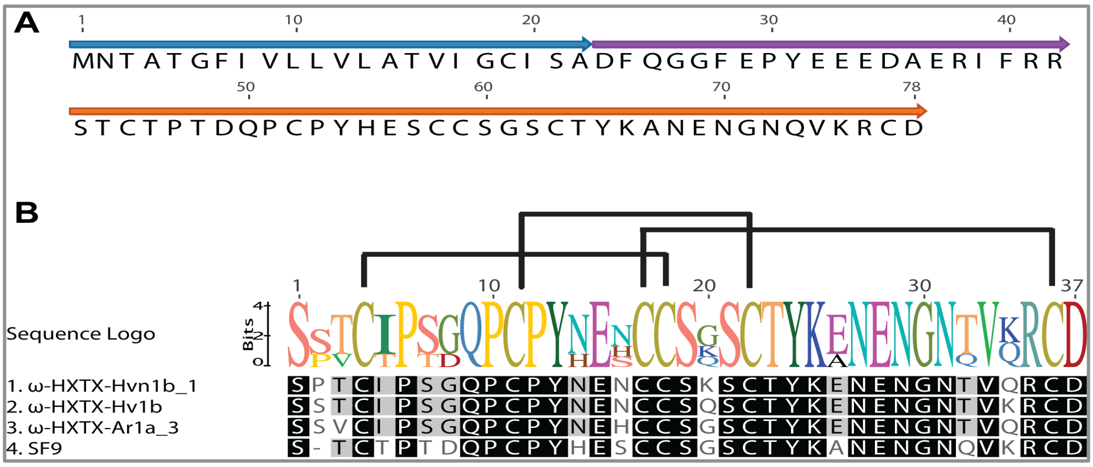


**Figure 1.9:** (a) Schematic of 78-residue precursor encoding a Superfamily 9 peptide. The signal peptide, propeptide, and mature toxin are shown in blue, purple, and orange, respectively. (b) Sequence alignment showing amino acid identities (boxed in black) between ω-hexatoxin-1 family members from the Australian funnel web spiders *H. infensa, H. venenata* (Hvn), *H. versuta*, and *A. robustus* (Ar); the disulfide bond pattern is shown above the sequence logo.

#### Superfamily 10 [Kisin]

Superfamily 10 corresponds to the previously described ω-hexatoxin-2 family. These peptides are expressed as 102-residue prepropeptides (Figure 1.10); the mature toxin comprises 42 residues with C-terminal amidation (not shown in Figure 1.10). The prototypic member of this superfamily, ω-HXTX-Hv2a, is the most potent blocker of insect $Ca_V$ channels discovered to date, with an $IC_{50}$ of 130 pM for inhibition of $Ca_V$ currents in bee brain neurons[19]. The 3D structure comprises a disulfide-rich region organized into a compact globular domain containing an ICK motif and a disordered C-terminal "tail" that is crucial for activity[19].

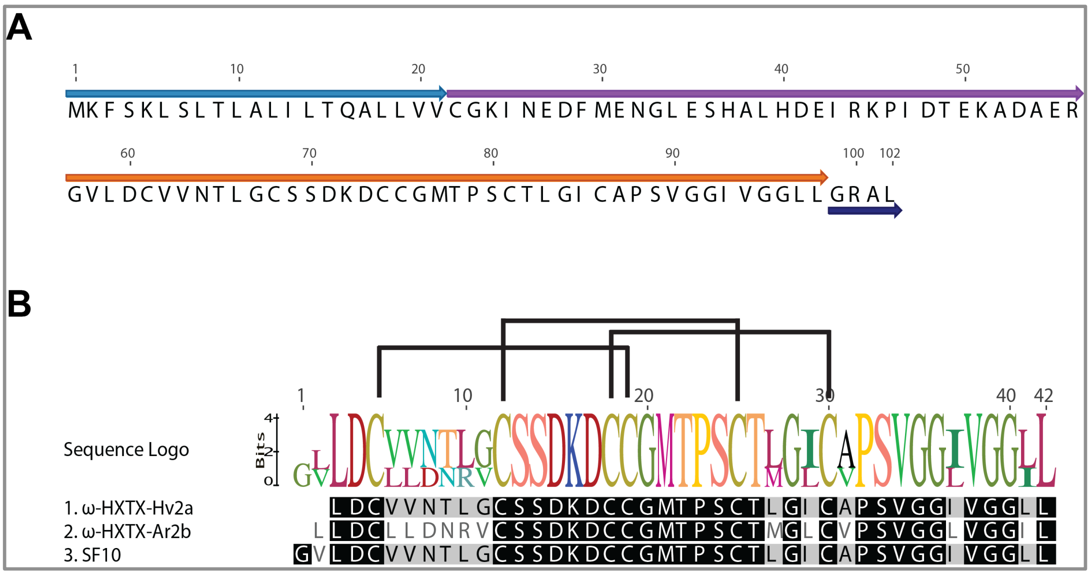


**Figure 1.10:** (a) Schematic of 102-residue precursor encoding the Superfamily 10 peptide ω-hexatoxin-Hi2a. The signal peptide, propeptide, mature toxin and C-terminal amidation signal are shown in light blue, purple, orange, and dark blue, respectively. (b) Sequence alignment showing amino acid identities (boxed in black) between members of the ω-hexatoxin-2 superfamily from *H. infensa*, *H. versuta*, and *A. robustus*; the disulfide-bond pattern shown above the sequence logo.

#### Superfamily 11 [Thanatos]

Superfamily 11 peptides (of which $U_7$-hexatoxin-Hi1a is the prototypic member; see Figure 1.11) are encoded by 105-residue precursors that lack a propeptide region. The mature toxin contains 82 residues. BLAST analysis revealed that this toxin is homologous to cystatins. Cystatins are a large superfamily of 120–140-residue peptides. They have been reported previously in the venoms of several organisms including snakes and spiders (i.e., the Chinese tarantula *Chilobrachys guanxiensis* and the black-widow spider *Latrodectus hesperus)* as well as in tick saliva.

Cystatins are currently divided into three families based on structural characteristics. Type 1 cystatins are cytoplasmic proteins comprising ~100 residues and lacking disulfide bridges. Type 2 cystatins are secreted peptides of ~120 residues with two disulfide bridges, whereas Type 3 cystatins comprise three repeats of the type 2 cystatins and encompass multiple domains[21]. Tick salivary cystatins display all the features described for Type 2 cystatins. It has been suggested that these cystatins might play a role in the regulation of endogenous cysteine peptidases involved in blood digestion and heme detoxification. Interestingly, the cystatin-like peptide isolated from *H. infensa* appears to be quite different to the cystatins isolated from the Chinese spider *C. jingzhao* and the saliva of the tick *Ornithodoros maubata* (Figure 1.11b). We predict that Superfamily 11 peptides represent a new scaffold for spider toxins and may also present a novel activity.

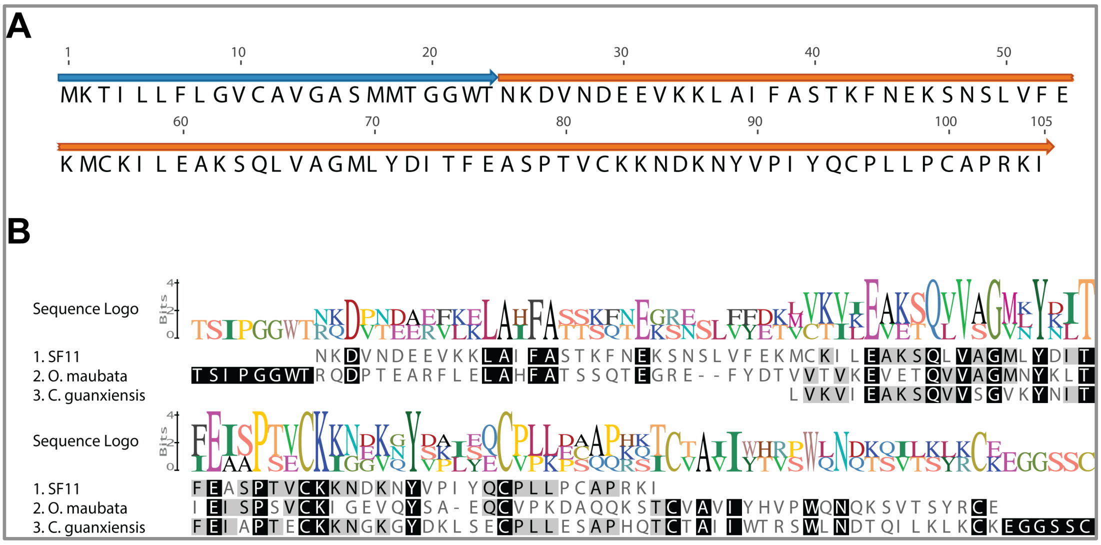


**Figure 1.11:** (a) Schematic of 105-residue precursor of the Superfamily 11 peptide $U_7$-hexatoxin-Hi1a. The signal peptide and mature toxin are shown in blue and orange, respectively. (b) Sequence alignment showing amino acid identities (boxed in black) between the *H. infensa* peptide and cystatins from the saliva of the tick *O. maubata* and the venom of the spider *C. guanxiensis*.

#### Superfamily 12 [Sekhmet]

Superfamily 12 peptides are expressed as 111-residue precursors that lack a propeptide region; the mature toxin spans 93 residues (Figure 1.12a). These peptides show homology to theraphotoxins from the Chilean rose tarantula *Grammostola rosea* and the barychelid spider *Tritamme loki.* These peptides also show weak homology to $U_{33}$-theraphotoxin-Cg1c and $U_{19}$-ctenitoxin-Pn1a as well as the MIT-like peptides found in Superfamily 2. However, the level of sequence identity between Superfamily 2 and Superfamily 12 peptides is only 23%, and the C-terminal region is significantly longer in the Superfamily 12 peptides (Figure 1.12b).

It is likely that Superfamily 12 and Superfamily 2 peptides evolved from the same ancestral gene, with sequence divergence and scaffold minimization in Superfamily 12 ultimately leading to a significant divergence between the two superfamilies. Whether the two families of toxins retain any functional similarity remains to be determined.

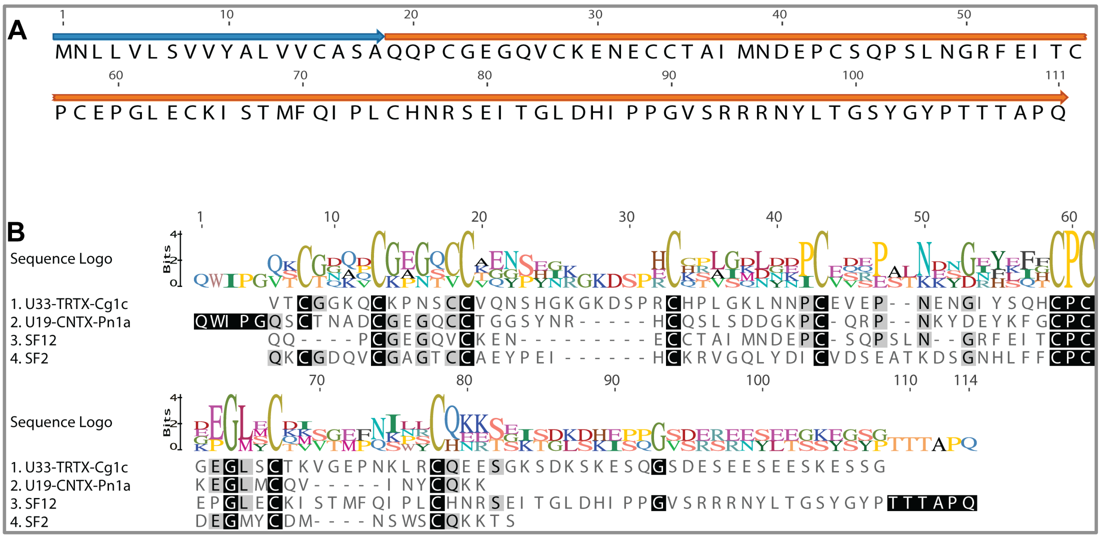

**Figure 1.12:** (a) Schematic of 111-residue precursor encoding Superfamily 12 peptides. The signal peptide and mature toxin are shown in blue and orange, respectively. (b) Sequence alignment showing amino acid identities (boxed in black) between $U_{33}$-theraphotoxin-Cg1c, $U_{19}$-ctenitoxin-Pn1a, $U_1$-hexatoxin-Hi1a (MIT-like peptide from Superfamily 2), and $U_8$-hexatoxin-Hi1a from Superfamily 12.

#### Superfamily 13 [Ares]

Superfamily 13 corresponds to the family of lethal δ-toxins from Australian funnel-web spiders. These peptides delay inactivation of $Na_V$ channels, resulting in prolonged action potentials that cause massive neurotransmitter release from both somatic and autonomic nerves[22,23]. The toxins are expressed as 115-residue prepropeptides, with the mature toxin comprising 40–42 residues (Figure 1.13). These toxins are stabilized by four disulfide bonds, three of which form an ICK motif[15,24].

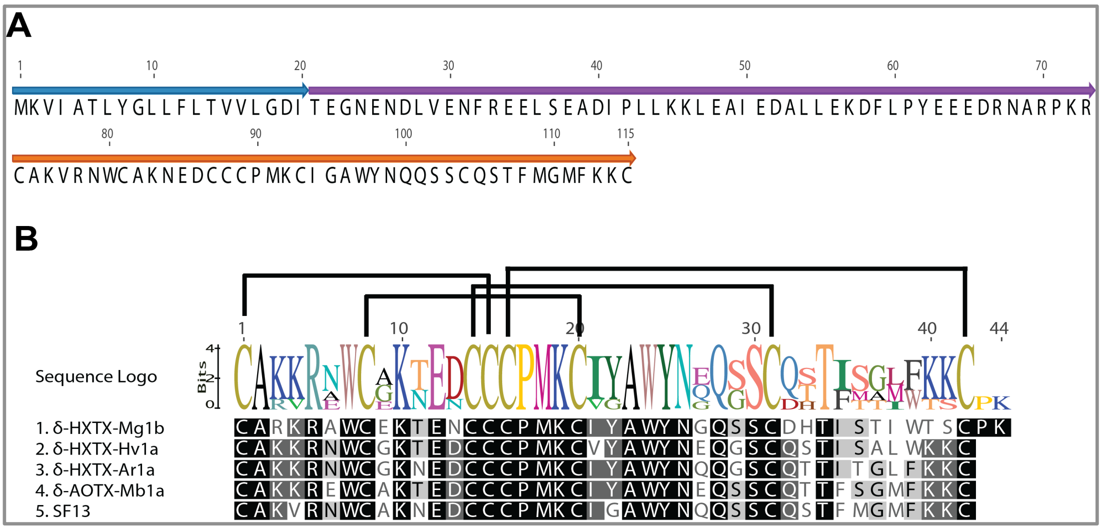

**Figure 1.13:** (a) Schematic of 115-residue precursor encoding the Superfamily 13 peptide δ-hexatoxin-Hi1a. The signal peptide, propeptide, and mature toxin are shown in blue, purple, and orange, respectively. (b) Sequence alignment showing amino acid identities (boxed in black) between δ-hexatoxins isolated from *H. versuta* (δ-HXTX-Hv1a), *A. robustus* (δ-HXTX-Ar1a), the Australian Eastern mouse spider Missulena bradleyi (δ-actinopoditoxin-Mb1a), and *Macrothele gigas* (δ-HXTX-

Mg1a/Magi-4); the disulfide-bond pattern shown above the sequence logo.

#### Superfamily 14 [Wurrukatte]

Superfamily 14 peptides are expressed as 119-residue prepropeptides, with the mature peptide spaning 69 residues (Figure 1.14a). These peptides show homology to toxins of unknown function and structure from the spiders *Trittame loki, Macrothele gigas*, and *C. guanxiensis* (Figure 1.14b). Interestingly, Superfamily 14 peptides have eight cysteine residues, with the last two separated by an unusually large gap of 28 residues. We therefore anticipate that these toxins will have a novel 3D fold.

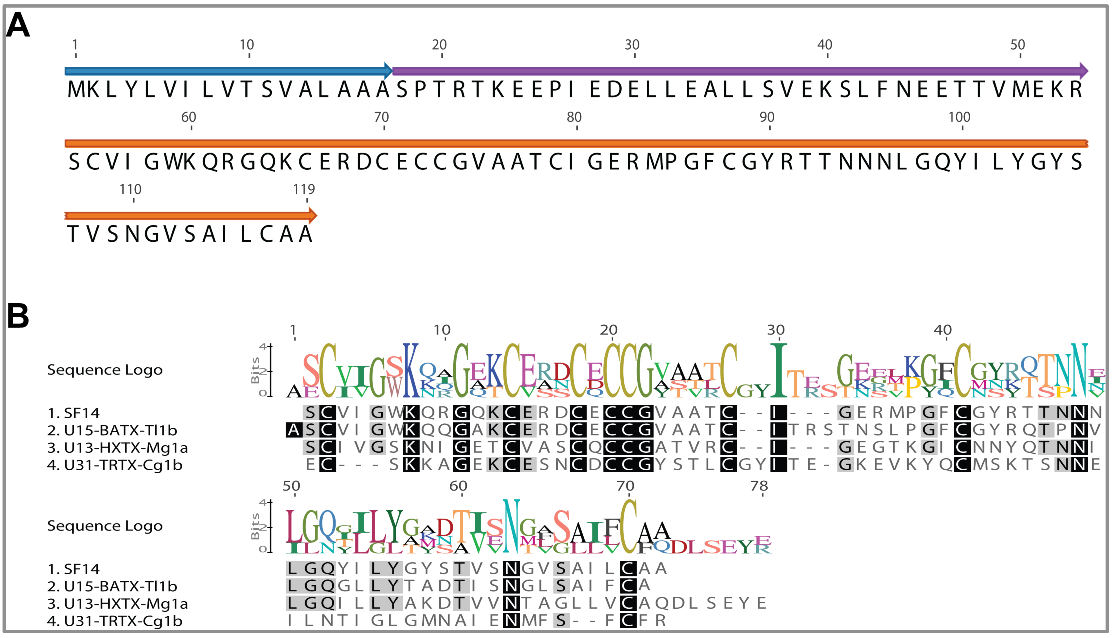


***Figure 1.14:*** (a) Schematic of 119-residue precursor encoding the Superfamily 14 peptide U$_9$-hexatoxin-Hi1a. The signal peptide, propeptide, and mature toxin are shown in blue, purple, and orange, respectively. (b) Sequence alignment showing amino acid identities (boxed in black) between the Superfamily 14 peptide U$_9$-hexatoxin-Hi1a and toxins from *T. loki*, *M. gigas* and *C. guanxiensis*.

#### Superfamily 15 [Pluto]

Superfamily 15 peptides are expressed as 132-residue prepropeptides. The mature peptide spans 101 amino acid residues and the 11-residue propeptide is unusually small, although it is acidic like most spider-toxin propeptides (Figure 1.15a). BLAST analysis revealed weak homology with an uncharacterised protein win the genome of the spider *Stegodypus mimosarum* but no other hits. Hence this appears to be a completely novel superfamily of venom peptides. These toxins contain 10 cysteine residues that are predicted to form five disulfide bonds. Although the most likely propeptide cleavage site is after the “VR” motif at positions 30–31, an alternative propeptide cleavage site is present after the “ER” motif at positions 58–59, as shown in Figure 1.15b. Proteomic analysis of the venom will be required to determine which is correct. However, regardless of the propeptide cleavage site, it is clear that this superfamily of venom peptides is unique.

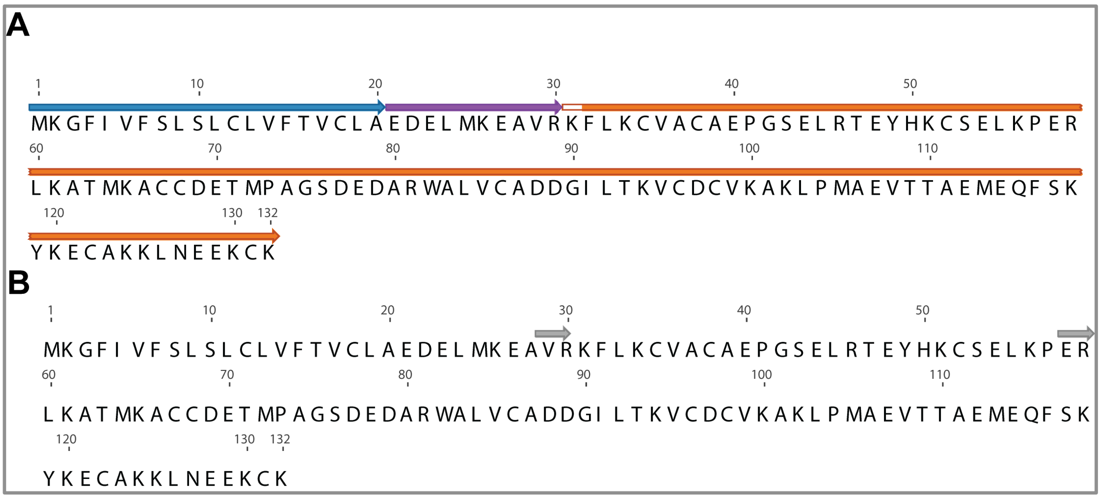


***Figure 1.15:*** *(a) Schematic of the 132-residue precursor encoding a highly novel Superfamily 15 peptide. The signal peptide, propeptide, and mature toxin are shown in blue, purple, and orange, respectively. (b) Toxin precursor showing alternative propeptide cleavage sites in gray.*

#### Superfamily 16 [Zahhak]

Superfamily 16 toxins are expressed as 71-residue precursors that comprise a signal peptide, propeptide, andmature toxin of 34 amino acid residues (Figure 1.16a). BLAST analysis revealed no sequence match in any database, but it is likely that this peptide will conform to an ICK fold.

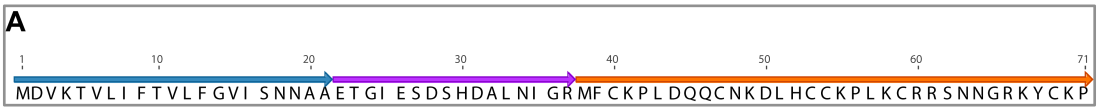


**Figure 1.16:** (a) Schematic of 71-residue precursor encoding the Superfamily 16 peptide $U_{11}$-hexatoxin-Hi1a. Signal peptide, propeptide and mature toxin are shown in blue, purple and orange, respectively.

#### Superfamily 17 [Nergal]

Superfamily 17 peptides are expressed as 115-residue prepropeptides precursors that are processed to yield a mature peptide of 44 amino acid residues (Figure 1.17a). BLAST analysis revealed that these peptides share homology with $U_{16}$-barytoxin-Tl1e, a peptides with unknown function and structure from the spider *Trittame loki*, and weak homology to the peptides μ-hexatoxin-Mg1c and ω-theraphotoxin-Asp2b isolated from the spiders *M. gigas* and *Aphonopelma sp.*, respectively (Figure 1.17b). Based on sequence homology with μ-hexatoxin-Mg1a, μ-hexatoxin-Mg1c is thought to be an insecticidal $Na_V$ channel modulator[25] whereas ω-theraphotoxin-Asp2b has been reported to inhibit $Ca_V$ channels in rat cerebellar granule cells (Patent WO 1994/010196 A1, 11/05/1994). Thus, the molecular target of Superfamily 17 peptides is difficult to predict.

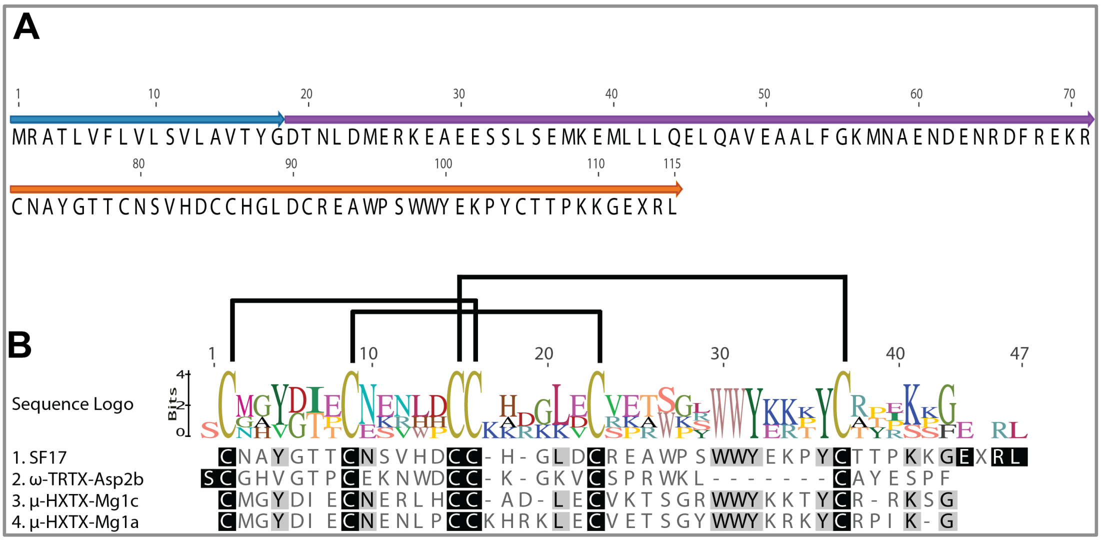


**Figure 4.22:** (a) Schematic of 115-residue precursor encoding the Superfamily 17 peptide $U_{12}$-hexatoxin-Hi1a. The signal peptide, propeptide, and mature toxin are shown in blue, purple, and orange, respectively. (b) Sequence alignment showing amino acid identities (boxed in black) between the Superfamily 17 member $U_{12}$-hexatoxin-Hi1a and μ-hexatoxin-Mg1c, μ-hexatoxin-Mg1a, and ω-theraphotoxin-Asp2b. Disulfide bonds are shown above the sequence logo.

#### Superfamily 18 [Grim Reaper]

Superfamily 18 toxins are expressed as 137-residues precursors without a propeptide region that are processed to yield an 87-residue mature toxin (Figure 1.18a). The C-terminal region of the prototypic family member $U_{13}$-hexatoxin-Hi1a is highly homologous to κ-theraphotoxin-Hs1a isolated from the venom of the Chinese bird spider *H. schmidti* but $U_{13}$-hexatoxin-Hi1a contains an additional 33-residue N-terminal region that includes four additional cysteine residues (Figure 1.18b). $U_{13}$-hexatoxin-Hi1a is also homologous to $U_1$-aranetoxin-Av1a that, unlike κ-theraphotoxin-Hs1a, also contains an extended N-terminal region (Figure 1.18b).

However, in contrast with $U_{13}$-hexatoxin-Hi1a, this extended N-terminal region does not contain additional cysteine residues. This raises an interesting evolutionary conundrum: is κ-theraphotoxin-Hh1a a minimized version of the more ancestral $U_{13}$-hexatoxin-Hi1a, or conversely is $U_{13}$-hexatoxin-Hi1a an extended version of the more ancestral κ-theraphotoxin-Hs1a? Information on other paralogs/orthologs will be required to answer this question. The structure of κ-theraphotoxin-Hs1a solved using NMR revealed a Kunitz-type fold possessing a C-terminal α-helix and a triple-stranded antiparallel β-sheet. The toxin is an unusual peptide in that it appears to have a dual function; it blocks vertebrate $K_V$ currents in rat dorsal root ganglion neurons (as well as $K_V1.1$ heterologously expressed in *Xenopus* oocytes) and it inhibits serine proteases such as trypsin. κ-theraphotoxin-Hs1a is lethal when injected into mouse brain with an $LD_{50}$ of 41.5 pmol/g (calculated based on literature value of 256 μg/kg and a toxin mass of 6166.05 Da)[26]. The high level of sequence identity between κ-theraphotoxin-Hs1a and the N-terminal region of $U_{13}$-hexatoxin-Hi1a suggests that Superfamily

18 peptides might also block $K_V$ channels and inhibit serine proteases. However, the four additional cysteine residues in the N-terminal extension in $U_{13}$-hexatoxin-Hi1a raises the possibility that the 3D fold and function of the *H. infensa* toxin might be completely different due to possession of a markedly different disulfide framework.

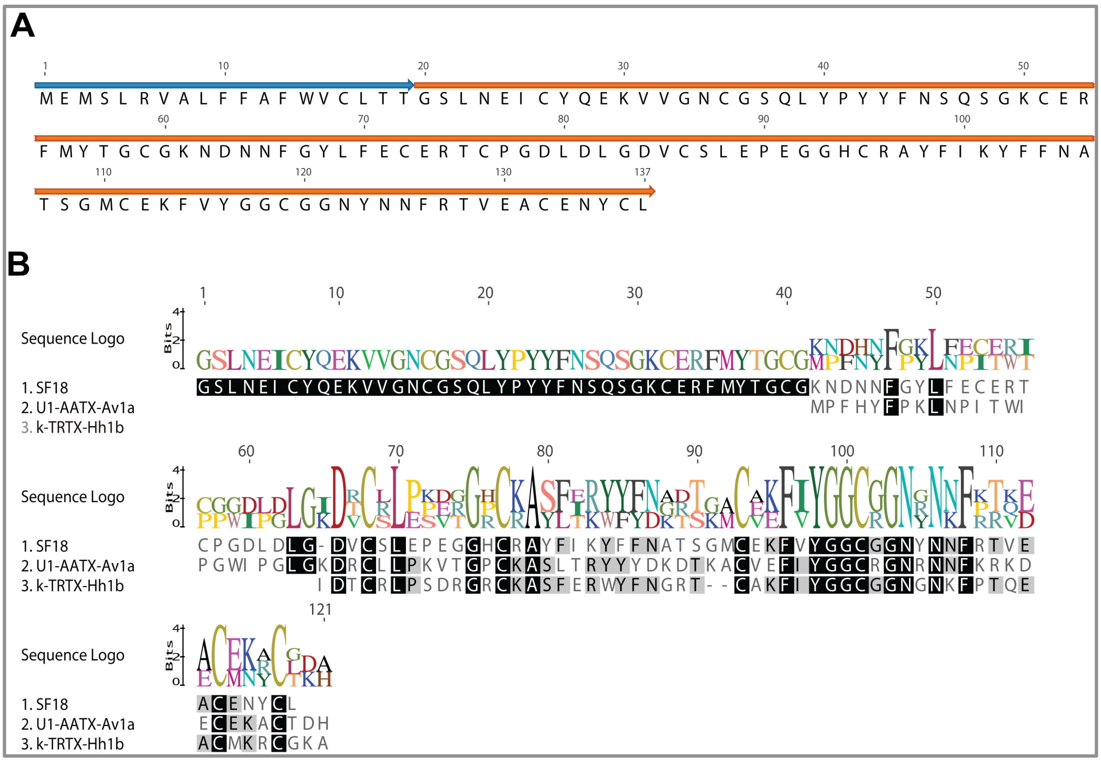


**Figure 1.18:** (a) Schematic of 137-residue precursor encoding the Superfamily 18 peptide $U_{13}$-hexatoxin-Hi1a. The signal peptide, propeptide, and mature toxin are shown in blue, purple, and orange, respectively. (b) Sequence alignment showing amino acid identities (boxed in black) between κ-theraphotoxin-Hs1a, $U_1$-aranetoxin-Av1a, and the Superfamily 18 peptide $U_{13}$-hexatoxin-Hi1a.

#### Superfamily 19 [Yama]

Superfamily 19 peptides are expressed as 76-residue precursors that are processed to yield a 58-residue mature toxin (Figure 1.19a). These peptides have no homology to known toxins in the database, and precursors of 6, 8 and 10 Cysteine bonds were found in the transcriptome analysis.

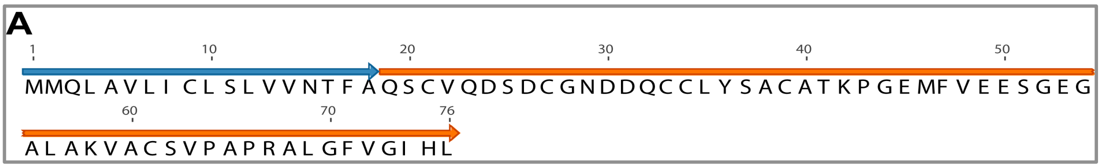


**Figure 1.19:** (a) Schematic of 76-residue precursor encoding the Superfamily 19 peptide $U_{14}$-hexatoxin-Hi1a. The signal peptide and mature toxin are shown in blue and orange, respectively.

#### Superfamily 20 [Ereshkigal]

Superfamily 20 peptides are expressed as 112-residue prepropeptide precursors that are processed to yield a mature peptide of 39 residues (Figure 1.20a). These peptides have homology with $U_{11}$-theraphotoxin-Hh1b, $U_{10}$-theraphotoxin-Hs1c, and ω-theraphotoxin-Bs2a isolated from venom of the spiders *H. hainanum*, *H. schmidti* and the Mexican red-knee spider *Brachypelma smithi,* respectively (Figure 1.20b). ω-theraphotoxin-Bs2a has modest insecticidal activity[27]; it weakly blocks insect $Na_V$ channels but has no effect on mammalian $Na_V$1.2, $Na_V$.14, $Na_V$1.5, and $Na_V$1.6 channels. Based on homology with ω-theraphotoxin-Asp3a from a related species (*Aphonopelma sp),* this family of toxins are thought to block $Ca_V$ channels[27].

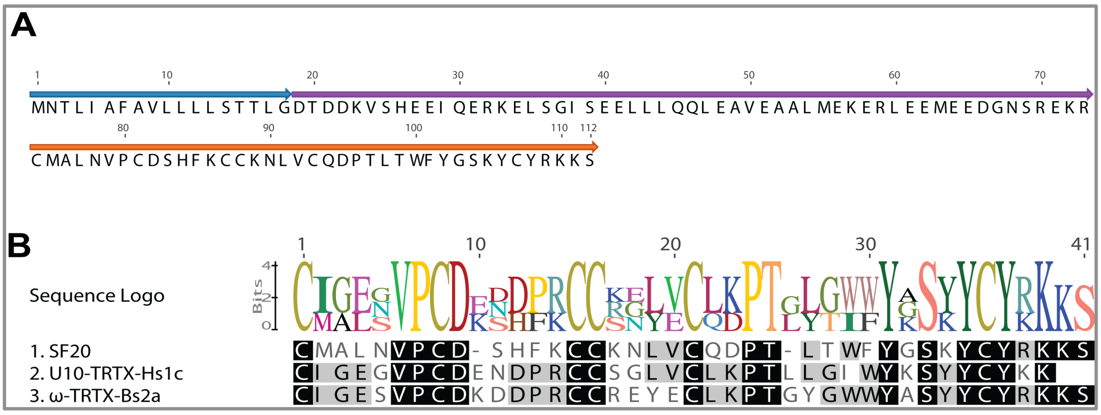


***Figure 1.20:*** *(a) Schematic of 112-residue precursor encoding the Superfamily 20 peptide $U_{15}$-hexatoxin-Hi1a. The signal peptide, propeptide, and mature toxin are shown in blue, purple, and orange, respectively. (b) Sequence alignment showing identities (boxed in black) between $U_{10}$-theraphotoxin-Hs1c, ω-theraphotoxin-Bs2a and $U_{15}$-hexatoxin-Hi1a.*

#### Superfamily 21 [Anubis]

Superfamily 21 peptides are expressed as 109-residue prepropeptides that are processed to yield a 66-residue mature peptide (Figure 1.21). Superfamily 21 peptides do not give sequence matches with any known toxins. Thus, this toxin superfamily is novel and with only four cysteine residues these peptides are likely to have a novel 3D fold.

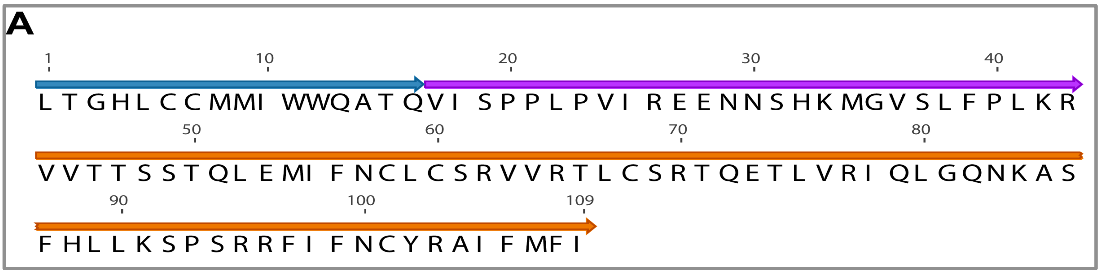


**Figure 1.21:** (a) Schematic of 109-residue precursor encoding the Superfamily 21 peptide $U_{16}$-hexatoxin-Hi1a. The signal peptide, propeptide, and mature toxin are shown in blue, purple, and orange, respectively.

#### Superfamily 22 [Loke]

Superfamily 22 peptides are expressed as 82-residue precursors that lack a propeptide and which are processed to yield a 63-residue mature toxin (Figure 1.22). The prototypic family member $U_{17}$-hexatoxin-Hi1a shows sequence homology with $U_{33}$-theraphotoxin-Cg1c from the venom of *C. guanxiensis*. The transcript encoding $U_{33}$-theraphotoxin-Cg1c has a very small propeptide (QEEEEP) and this has been entirely discarded in the transcript encoding the *H. infensa* toxin. The size of several of the intercystine loops has also been reduced in $U_{17}$-hexatoxin-Hi1a.

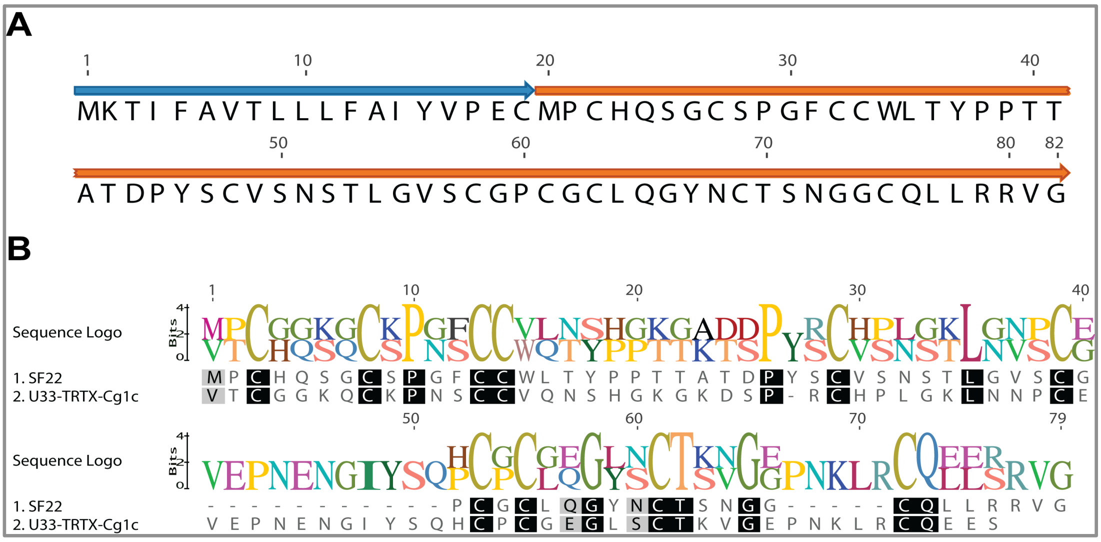


**Figure 1.22:** (a) Schematic of 82-residue precursor encoding the superfamily 22 peptide $U_{17}$-hexatoxin-Hi1a. The signal peptide and mature toxin are shown in blue and orange, respectively. (b) Sequence alignment showing amino acid identities (boxed in black) between $U_{33}$-theraphotoxin-Cg1c and $U_{17}$-hexatoxin-Hi1a.

#### Superfamily 23 [Qamaits]

Superfamily 23 peptides are expressed as ~88-residue prepropeptides that are processed to yield a mature peptide comprising 40–60 amino acid residues (Figure 1.23a). These peptides have weak homology with $U_6$-lycotoxin-Ls1c, δ-amaurobitoxin-Pl1c and $U_5$-pisautoxin-Dm1a isolated from venom of the spiders *Lycosa singoriensis*, *Pireneitega luctuosa* and *Dolomedes mizhoanus*, respectively (Figure 1.23b). δ-Amaurobitoxin-Pl1c delays inactivation of insect $Na_V$ channels and is lethal to insects but not mice[28].

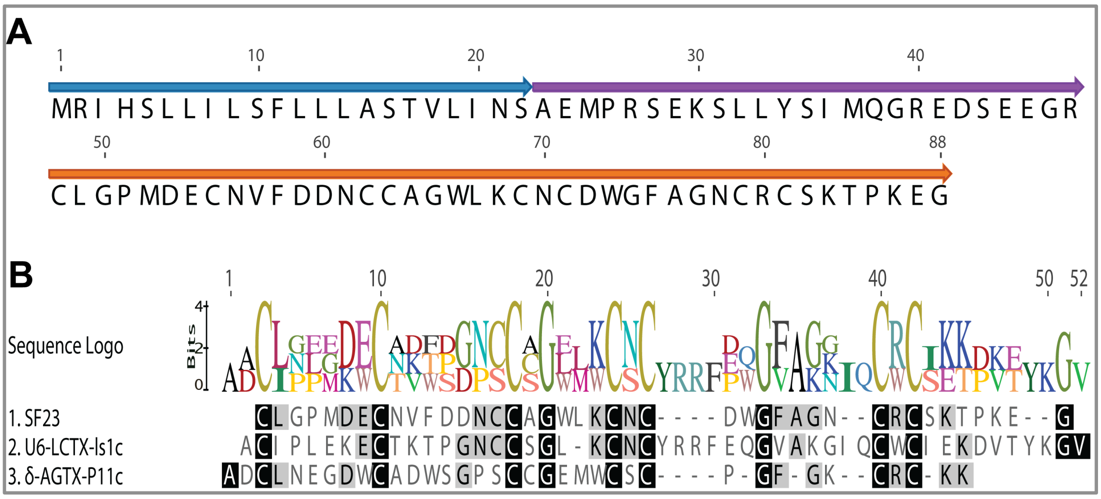


**Figure 1.23:** (a) Schematic of 88-residue precursor encoding the Superfamily 23 peptide $U_{18}$-hexatoxin-Hi1a. The signal peptide, propeptide, and mature toxin are shown in blue, purple, and orange, respectively. (b) Sequence alignment showing amino acid identities (boxed in black) between $U_6$-lycotoxin-Ls1c, δ-amaurobitoxin-Pl1c, and the Superfamily 23 peptide $U_{18}$-hexatoxin-Hi1a.

#### Superfamily 24 [Ankou]

Superfamily 24 peptides are expressed as 110-residue prepropeptide precursors. The mature toxin spans 39 residues and includes six cysteines that are predicted to form three disulfide bonds (Figure 1.24). The mature toxin has ~50% sequence identity with ω-TRTX-Bs2a, an ICK toxin from the Mexican red-knee tarantula *Brachypelma smithi.* ω-TRTX-Bs2a is a weak insecticidal toxin with modest effects on insect Nav channels, however based on homology with ω-TRTX-Asp3a from the spider *Aphonopelma sp.*, this toxin might also inhibit voltage-gated calcium channels.

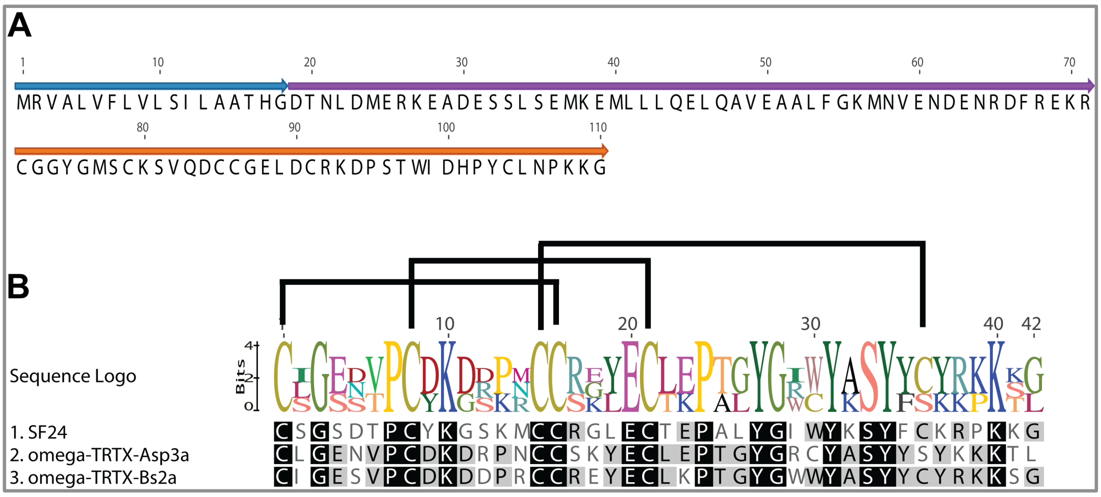


**Figure 1.24:** (a) Schematic of 110-residue precursor encoding the Superfamily 24 peptide $U_{19}$-hexatoxin-Hi1a. The signal peptide, propeptide, and mature toxin are shown in blue, purple, and orange, respectively. (b) Sequence alignment showing amino acid identities (boxed in black) between ω-TRTX-Bs2a, ω-TRTX-Asp3a and $U_{19}$-hexatoxin-Hi1a. Disulfide-bond patterns is shown above the sequence logo.

#### Superfamily 25 [Tezcatlipoca]

Superfamily 25 peptides are expressed as 145-residue precursors that lack a propeptide region and which are processed to yield an unusually large mature toxin comprising ~130 amino acid residues (Figure 1.25a). The prototypic family member $U_{20}$-hexatoxin-Hi1a appears to be an ortholog of $U_{24}$-ctenitoxin-Pn1a isolated from the venom of the highly venomous Brazilian armed spider *Phoneutria nigriventer* (Figure 1.25b) and the more recently discovered peptide LTDF s-18 from the fishing spider *Dolomedes fimbriatus*[29].

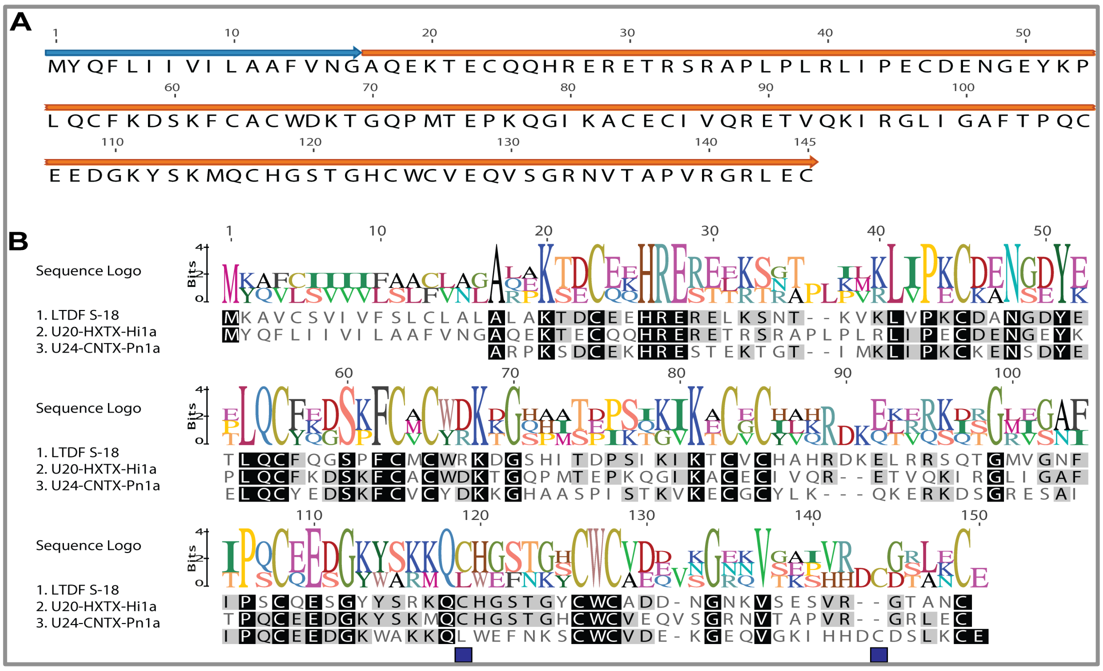


**Figure 1.25:** (a) Schematic of 145-residue precursor encoding the Superfamily 25 peptide $U_{20}$-hexatoxin-Hi1a. The signal peptide and mature toxin are shown in blue and orange, respectively. (b) Sequence alignment showing amino acid identities (boxed in black) between $U_{24}$-ctenitoxin-Pn1a, LTDF s-18, and $U_{20}$-hexatoxin-Hi1a. Unmatched cysteine residues are highlighted with purple boxes.

$U_{24}$-ctenitoxin-Pn1a is an unusual peptide that contains 12 cysteine residues and has weak homology to cysteine proteinase inhibitors. It contains two thyroglobulin type 1 domains mapped to residues 4–67 and 72–127. Disulfide bonds for eight of the cysteine residues have been predicted based on homology to these inhibitors. The sequence position of 11 of the 12-cysteine residues is conserved in the *H. infensa* toxin (Figure 1.25b) and therefore its structure and function may be similar to the toxins from *Phoneutria* and *Dolomedes*. Few peptides of this kind have been described in spider venoms primarily because its mass lies outside the typical range of 1–10 kDa sampled in most mass spectrometric studies of spider venoms. Thus, Superfamily 25 peptides represent an interesting challenge for future structure-function studies.

#### Superfamily 26 [Set]

Superfamily 26 toxins are expressed as 92-residue precursors that comprise a signal peptide and a mature toxin of 73 amino acid residues, but lack a propeptide region (Figure 1.26a). These peptides appear to be orthologs of the potent insecticidal toxin $U_1$-CUTX-As1a from venom of the trapdoor spider *Apomastus* *schlingeri* (Figure 1.26b). $U_1$-CUTX-As1a induces rapid, irreversible flaccid paralysis in the lepidopteran *Manduca sexta* that ultimately results in death with an $LD_{50}$ of 174 pmol/g[30]. Identity between $U_1$-CUTX-As1a and SF26 is ~61% (42 identical residues) with both toxins containing a long N-terminal "tail" preceding the first cysteine residue. Preliminary results showed that $U_{21}$-hexatoxin-Hi1a induces reversible paralysis in blowflies (*L. cuprina*) with a $PD_{50}$ after 24 h of 153 ± 23 pmol/g (unpublished results).

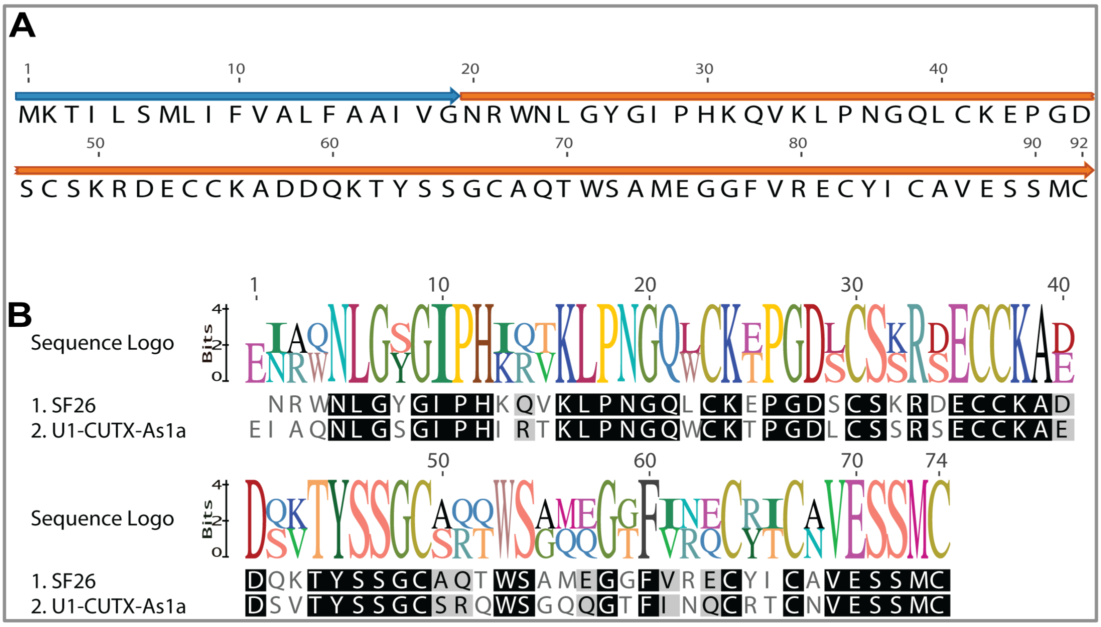


**Figure 1.26:** (a) Schematic of 92-residue precursor encoding the Superfamily 26 peptide $U_{21}$-hexatoxin-Hi1a. The signal peptide and mature toxin are shown in blue and orange, respectively. (b) Sequence alignment showing amino acid identities (boxed in black) between $U_1$-CUTX-As1a from *A. schilingeri* and the Superfamily 26 peptide.

#### Proteins, enzymes and secreted peptides

##### Superfamily 27 [Mara]

Superfamily 27 peptides are expressed as prepropeptide precursors of ~167 residues that are processed to yield a 132-residue mature peptide (Figure 1.27a). BLAST searches revealed moderate homology (29–35% identity) with a variety of non-venom triacylglycerol lipases (TGL) from the deer tick *Ixodes scapularis*, the jumping ant *Harpegnathos saltator*, the honeybee *Apis mellifera*, and various other insects as well as a putative lipase transcript found in the sialotranscriptome of the Gulf Coast tick *Amblyomma maculatum* (AEO33734.1). However, the best match (46% identity) is with the partial sequence of a venom-gland transcript from the King baboon spider *Pelinobius muticus* (UniProt D5J706)[31]. Even though transcripts encoding this class of putative lipases have now been found in tick salivary glands and spider venom glands, it remains to be shown unequivocally that they are expressed in spider venoms. We believe that this is likely as lipase activity has been reported in the venom of the endoparasitic wasps *Pimpla hypochondriaca* and *Chelonus inanitus* and the ectoparasitic wasp *Nassonia vitripennis*[32-34] and the Superfamily 30 proteins show significant homology with these lipases (Figure 1.27b). Triacylglycerol lipases are lipolytic enzymes that hydrolyse the ester linkage of triglycerides to yield monoglycerides and free fatty acids. It is unclear what function this activity might play in spider venom, other than possibly a predigestive role.

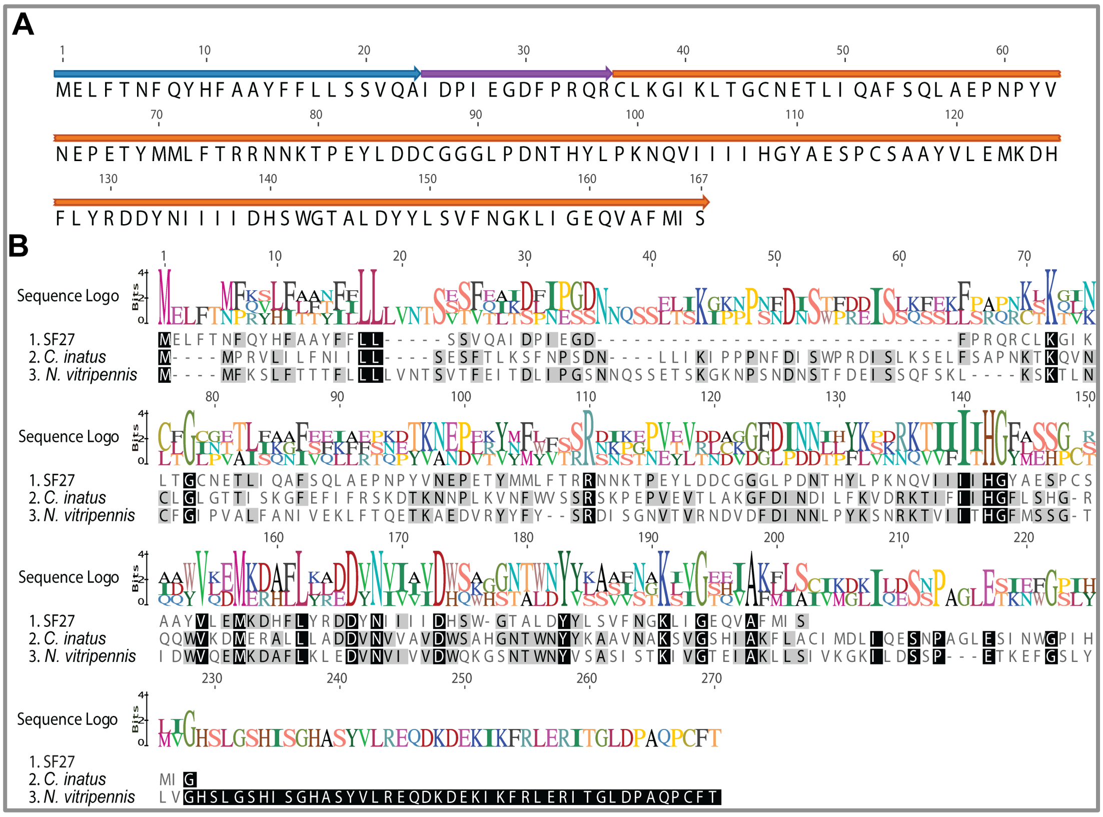

**Figure 1.27:** (a) Schematic of 165-residue precursor encoding Superfamily 27 lipases. The signal peptide and mature toxin are shown in blue and orange, respectively. (b) Sequence alignment showing identities (boxed in black), between lipases from the venom of *H. infensa* and the parasitic wasps *Chelonus inanitus* and *Nasonia vitripennis* (truncated sequence).

#### Superfamily 28 [Tuoni]

Superfamily 28 peptides are expressed as 442-residue precursors that lack a propeptide region (Figure 1.28a). The precursor is processed to yield a large protein of ~425 residues that is homologous with hyaluronidases from a variety of animal venoms, including bees, cone snails, fish, lizards, scorpions, snakes, and wasps[35-41]. SF27 shows ~30% identity with hyaluronidases from bee and scorpion venoms (Figure 1.27b). Hyaluronidases hydrolyse hyaluronic acid (HA), a key component of the extracellular matrix of vertebrates[45]. It has been proposed that venom hyaluronidases act as "spreading factors" to facilitate dispersion of venom toxins so they have better access to their molecular target(s)[38,42-44].

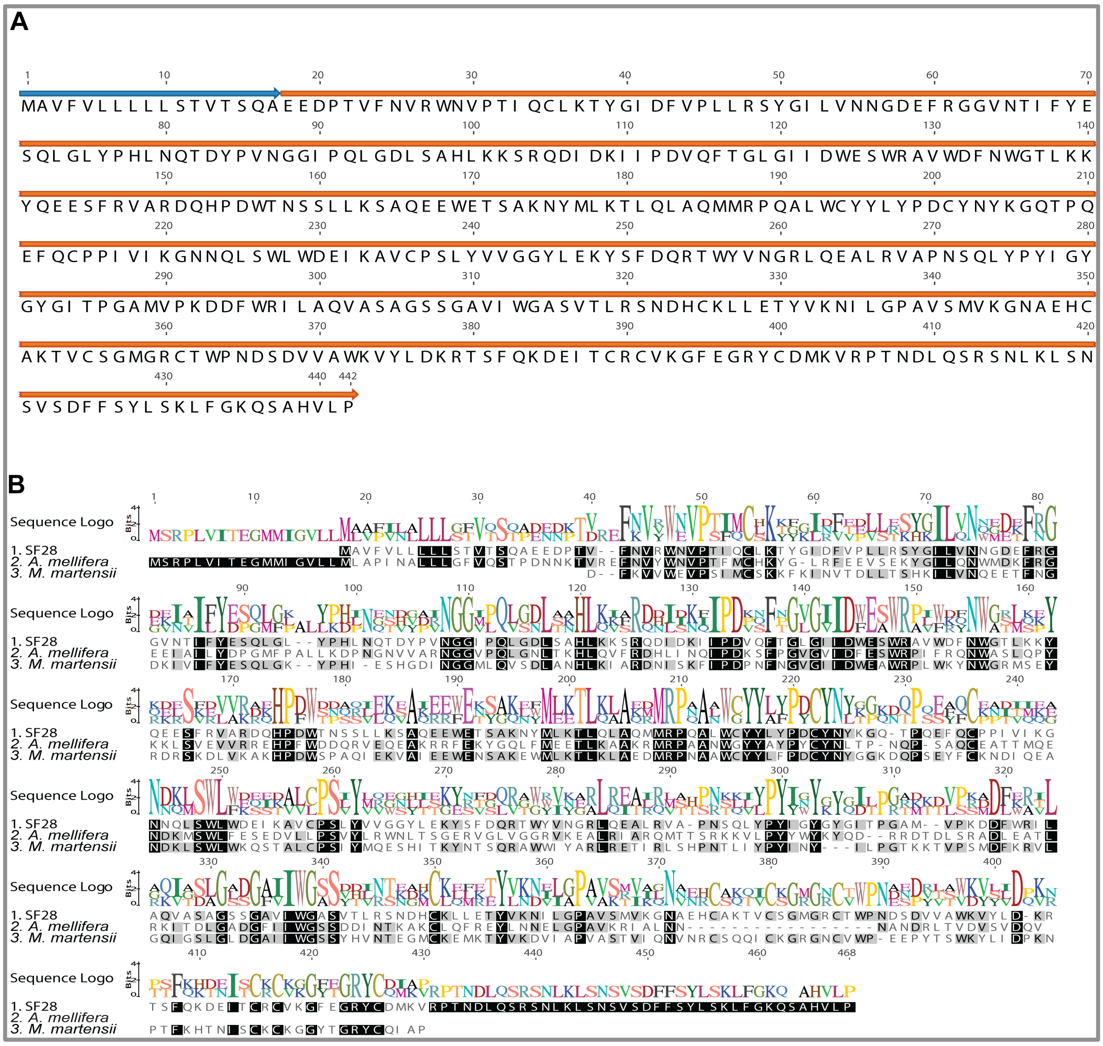

***Figure 1.28:*** (a) Schematic of 442-residue precursor encoding spider-venom hyaluronidase (Superfamily 28). The signal peptide and mature toxin are shown in blue and orange, respectively. (b) Sequence alignment showing amino acid identities (boxed in black) between venom hyaluronidases from the scorpion *Mesobuthus martensii*, the honeybee *Apis mellifera*, and the *spider H. infensa.*

#### Superfamily 29 [Mars]

Superfamily 29 proteins are expressed as precursors of >156 residues and only incomplete transcripts were obtained that were missing the N-terminal region of the mature toxin. Nevertheless, BLAST searches revealed strong homology between Superfamily 29 proteins and phospholipase $A_2$ ($PLA_2$) found in hymenopteran venoms (Figure 1.29). The best matches were with $PLA_2$ enzymes from the silk moth *Bombyx mori* (45% identity) and venom of the bumblebees *Bombus ignitus* (43% identity over 137 residues; UniProt C7B2L3), *Megabombus pennsylvanicus* (43% identity over 131 residues; UniProt Q7M416), and *Bombus terrestris* (40% identity over 137 residues; UniProt P82917). Moderate matches with 39% identity were also found with venom $PLA_2$ from the honeybees *Apis mellifera* (UniProt P00630) and *Apis cerana cerana* (UniProt Q9BMK4). $PLA_2$ are enzymes that catalyse the hydrolysis of phospholipids at the *sn*-2 bond of the glycerol backbone resulting in the release of lysophospholipids and fatty acids. They have a variety of derived functions in reptile venoms including antiplatelet, myotoxic, and neurotoxic activity[46] but their role in arthropod venoms is less well understood. There are at least 17 classes of secreted $PLA_2$s, but only groups IA, IIA, IIB, III, IX, and XII have been recruited into reptile venoms[46].

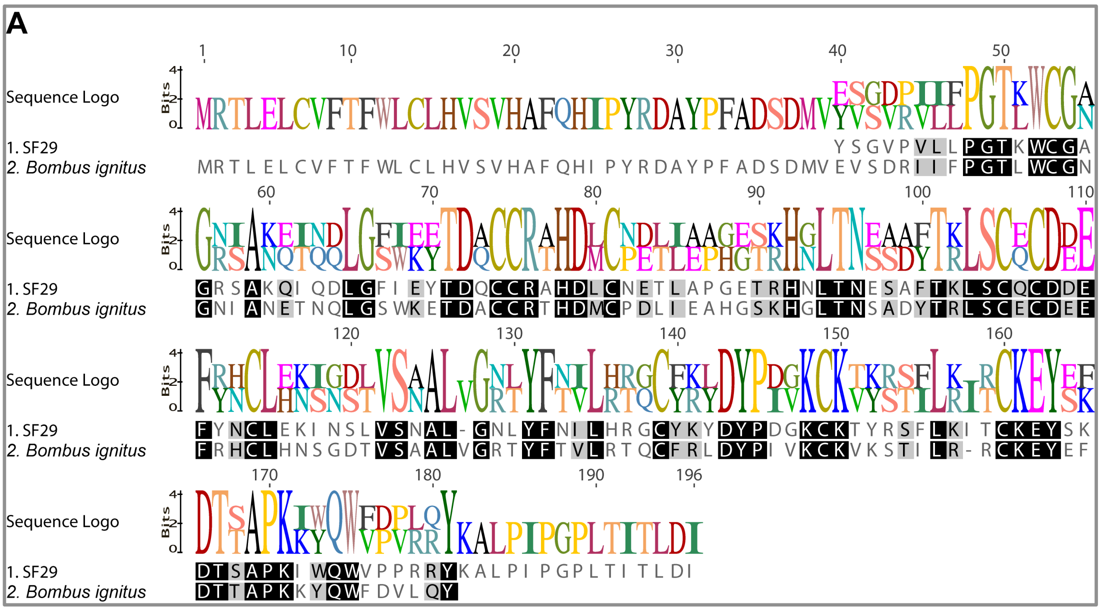


**Figure 1.29:** (a) Alignment of Superfamily 29 $PLA_2$ with $PLA_2$ isolated from the venom of the bumblebee *Bombus ignitus*. Amino acid identities are boxed in black.

#### Superfamily 30 [Sauron]

Superfamily 30 proteins are expressed as 368-residue prepropeptide precursors that are processed to yield large mature toxins (Figure 1.30a) that are homologous to invertebrate venom cysteine-rich secretory proteins (CRISPs) commonly known as venom allergen 5 (VA5) (Figure 1.30b). CRISPs are a large group of secreted proteins with molecular masses that range from 20 to 30 kDa. In mammals they are primarily as expressed in spermatocytes and granules of neutrophils and play roles in sperm maturation and host defense[47-50]. However, related proteins have been recovered in the venom or salivary secretions of stinging insects (bees, wasps, bugs, mosquitoes and ticks)[51-53], cephalopods[54], cone snails[55] and spiders[56,57]. These latter CRISPs have been shown to inhibit a variety of ion channels, including cyclic nucleotide-gated channels, ryanodine receptors, and calcium and potassium channels[46,58]. Notably, the CRISPs in arthropod venoms lack the C-terminal cysteine-rich domain contained in their metazoan counterparts[46].

The closest match to the *H. infensa* CRISP is a CRISP isolated from the Chilean rose tarantula *Grammostola rosea* (66.5% identity; UniProt M5AWW7), a CRISP/VA5 identified in a venom-gland transcriptome from the trapdoor spider *Trittame loki* (61.8% identity, UniProt W4VS53), a CRISP/VA5 from the velvet spider *Stegodyphus mimosarum* (37.4% identity, ENA A0A08UF27) and a CRISP identified in the sialo-transcriptome of the tick *Amblyomma maculatum* (31% identity over 224 residues; UniProt G3MIH5). Recently, the structure of the CRISP/VA5 protein Ves v5 from the blue jacket wasp *Vespula vulgaris* was solved using X-ray crystallography at 1.9 Å resolution. The structure is an α-β-α sandwich. The upper layer consists of three α-helices, two of them running perpendicular to the β-sheet and one running in parallel, while the β-sheet is composed of four antiparallel β strands. The lower layer comprises four α-helices [59]. Alignments of several CRISP/VA5 sequences isolated from spiders and hymenopteran venoms show that they share at least 30% identity with conservation of the important cysteine residues; thus the structure of the *H. infensa* CRISP is likely to resemble that of wasp Ves v 5. However, the function of CRISPs in the venom of *H. infensa* and other spiders remains enigmatic, and this should be a fruitful area for future investigations.

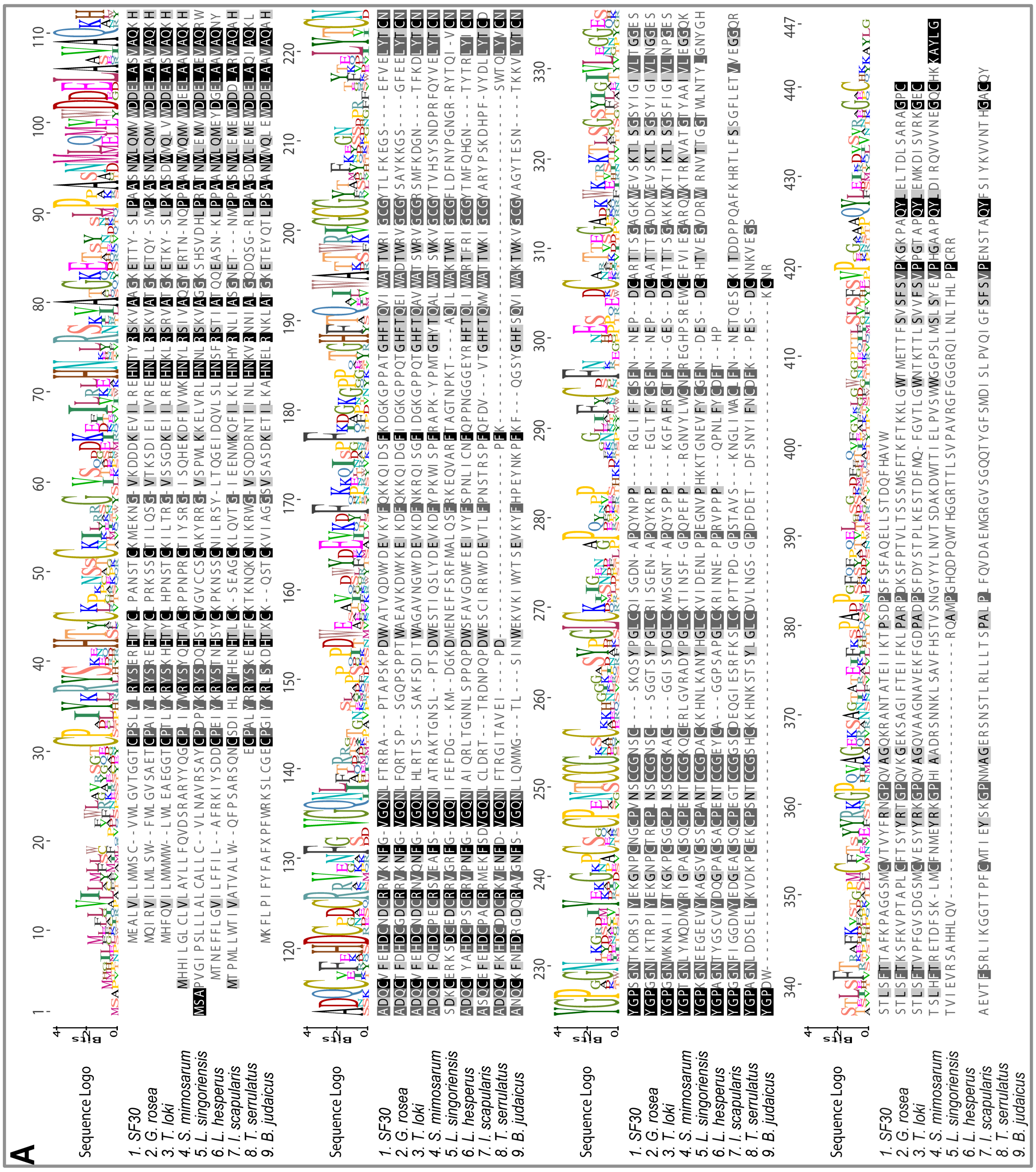

**Figure 1.30:** *H. infensa* CRISP aligned with CRISPs and venom allergens isolated from spiders (*G. rosea*, *T. loki*, *S. mimosarum*, *L. singoriensis* and *L. hesperus*), ticks (*Ixodes scapularis*), and scorpions (*Tityus serrulatus* and *Buthus judaicus*). Amino acid identities are boxed in black.

### Secreted proteins

#### Superfamily 31 [Hulda]

Superfamily 31 proteins are expressed as 180-residue precursors that lack a propeptide and which are processed to yield a 155-residue mature protein (Figure 1.31a). A BLAST search revealed homology between the prototypic Superfamily 31 protein (SF31) and venom-gland transcripts from the tarantula *Chilobrachys guanxiensis* and the brown huntsman spider *Heteropoda venatoria*. Transcriptomic analysis of the *C. guanxiensis* revealed three low abundance SF31 orthologs (JZTX-81, JZTX-82 and JZTX-83) that did not have homology with any known sequences. These proteins are highly enriched in cysteine residues, with 12 being present in JZTX-82, which is the closest match to Superfamily 31 (Figure 1.31b). The function of these novel venom proteins remains to be determined.

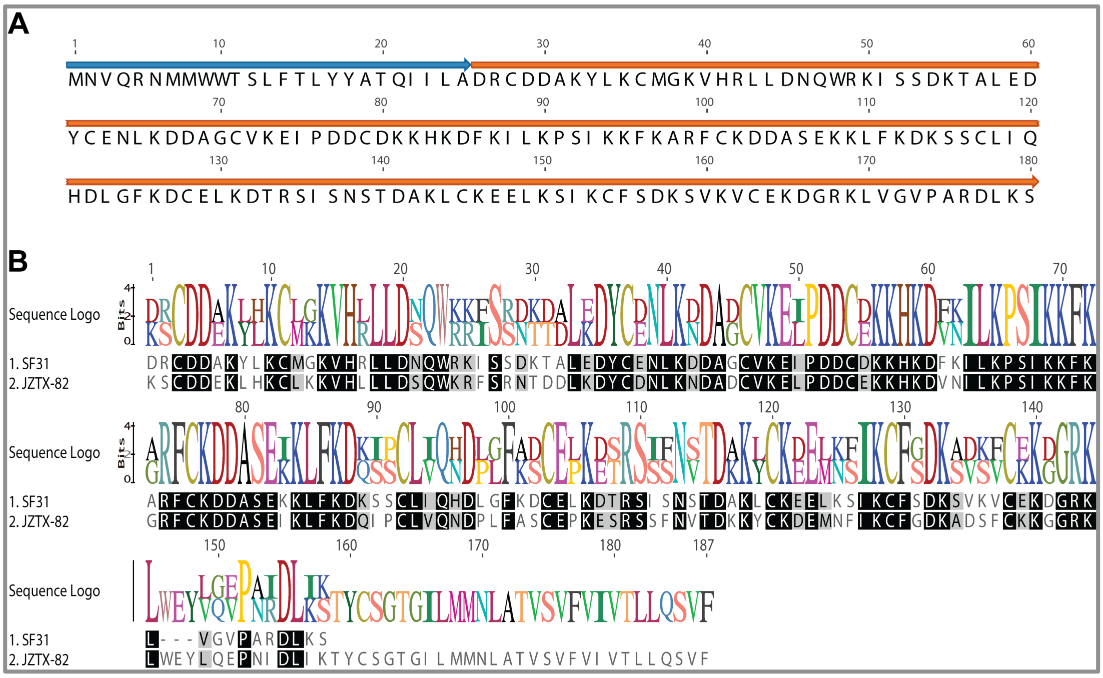

**Figure 1.31:** (a) Schematic of 180-residue precursor encoding a Superfamily 31 peptide. The signal peptide and mature toxin are shown in blue and orange, respectively. (b) Sequence alignment showing amino acid identities (boxed in black) between JZTX-82 and a Superfamily 31 peptide.

#### Superfamily 32 [Midgardsormen]

Superfamily 32 proteins are expressed as 220-residue precursors that comprise a signal peptide of 18 residues and a large mature peptide with 12 cysteines (Figure 1.32a). BLAST analysis of this superfamily revealed sequence homology to transcripts previously isolated from venom-gland cDNA libraries from the spiders *Chilobrachys guanxiensis and Pelinobius muticus* (Figure 1.32b). The function of these unusual venom proteins remains to be determined.

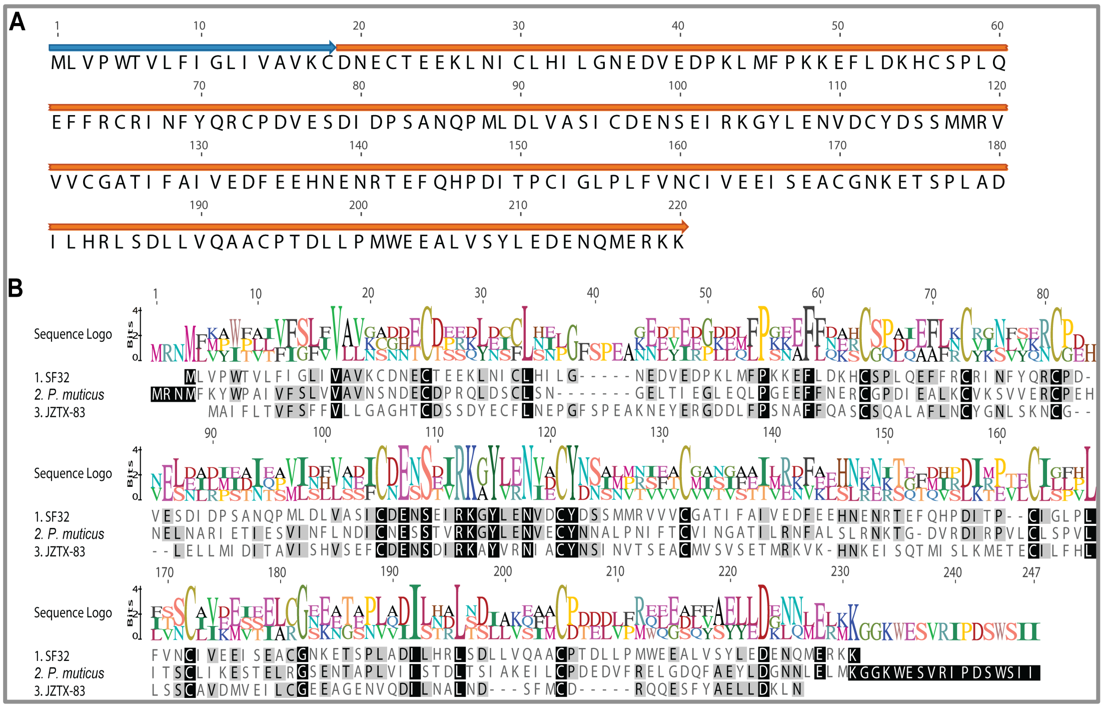

**Figure 1.32:** (a) Schematic of 220-residue precursor of a Superfamily 32 protein, with the signal peptide and mature protein highlighted in blue and orange, respectively. (b) Sequence alignment showing amino acid identities (boxed in black) between the Superfamily 32 protein, JZTX-83, and an ortholog in the spider *Pelinobius muticus.*

#### Superfamily 33 [Menhit]

Superfamily 33 proteins are expressed as precursors of more than 180 residues. BLAST analysis of the incomplete transcript obtained for this superfamily revealed sequence homology to lectins from horseshoe crabs, including techylectin-5A (UniProt Q9U8W8; PDB 1JC9) from *Tachypleus tridentatus* and carcinolectin 5a (UniProt A1KYQ0) from *Carcinoscorpius rotundicauda*. In invertebrates, the lack of an adaptive immune system has led to the development of various defense mechanisms that make up their so-called “innate immunity”, which enables recognition of invading pathogens via common antigens on their surface[60].

A well-characterized organism with such a defense mechanism is the Japanese horseshoe crab *T. tridentatus.* The horseshoe crab relies completely on innate immunity to defend itself from invading pathogens by producing lectin-type molecules in its hemocytes. To date, a total of five techylectins have been described and they all agglutinate human erythrocytes as well as both Gram-positive and Gram-negative bacteria. Hence, as for gomesin (Superfamily 5), Superfamily 33 appears to represent yet another example of spiders recruiting molecules usually associated with innate immunity into their venom. The role of the lectin-like Superfamily 33 proteins remains to be determined but, as discussed above for gomesin, it is unlikely they play a role in protecting the venom gland against invading pathogens. They more likely have a derived function. C-type lectins in snake venoms and the stinging bristles of the *Lonomia* caterpillar act on the vertebrate coagulation pathway, those from stonefish venom are myotoxic, and the lectins present in the feeding secretions of hematophagous insects are thought to have an antihemostatic function[46]. Thus, there are a variety of possible derived functions for the lectins in *H. infensa* venom.

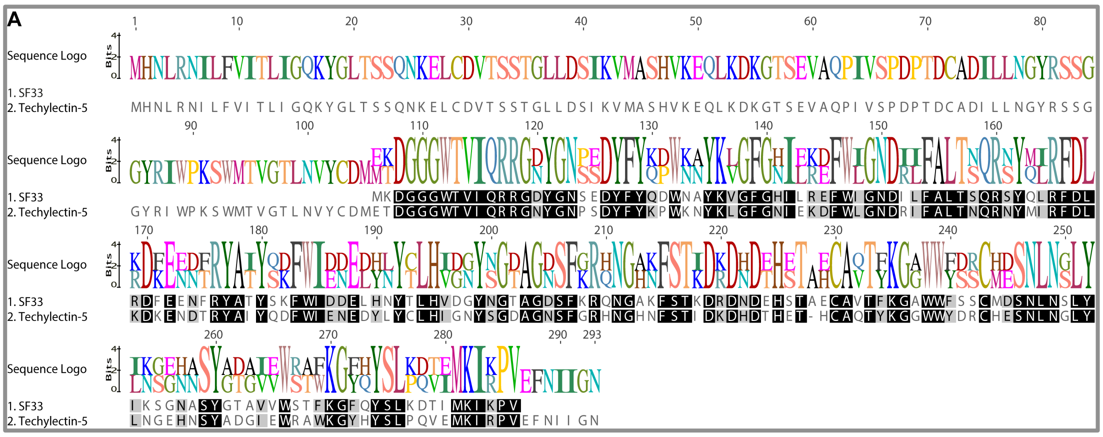


**Figure 1.33:** (a) Sequence alignment showing amino acid identities (boxed in black) between the Superfamily 33 protein (SF33) and techylectin 5 from a horseshoe crab.
